# The evolutionary history of mammoth wasps (Hymenoptera: Scoliidae)

**DOI:** 10.1101/2022.01.24.474473

**Authors:** Z. Khouri, J.P. Gillung, L.S. Kimsey

**Affiliations:** Bohart Museum of Entomology, University of California, Davis, CA, U.S.A; Lyman Entomological Museum, McGill University, Montreal, Quebec, Canada

## Abstract

Scoliid wasps comprise a clade of aculeate insects whose larvae are parasitoids of scarabaeid beetle grubs. While scoliids have been studied and used as biological control agents, research into the group’s evolution, as well as the stability of scoliid taxonomy, has been limited by a lack of reliable phylogenies. We use ultraconserved element (UCE) data under concatenation and the multispecies coalescent to infer a phylogeny of the Scoliidae. In order to mitigate potential issues arising from model misspecification, we perform data filtering experiments using posterior predictive checks and matched-pairs tests of symmetry. Our analyses confirm the position of *Proscolia* as sister to all other extant scoliids. We also find strong support for a sister group relationship between the campsomerine genus *Colpa* and the Scoliini, rendering the Campsomerini non-monophyletic. Campsomerini excluding *Colpa* (hereafter Campsomerini *sensu stricto*) is inferred to be monophyletic, with the Australasian genus *Trisciloa* recovered as sister to the remaining members of the group. Out of nine genera in which more than one species was sampled, *Campsomeriella*, *Dielis*, *Megascolia*, and *Scolia* are inferred to be non-monophyletic. Analyses incorporating fossil data indicate an Early Cretaceous origin of the crown Scoliidae, with the split between Scoliini + *Colpa* and Campsomerini *s.s.* most probably occurring in the Late Cretaceous. Posterior means of Scoliini + *Colpa* and Campsomerini *s.s.* crown ages are estimated to be in the Paleogene, though age 95% HPD intervals extend slightly back past the K-Pg boundary, and analyses including fossils of less certain placement result in more posterior mass on older ages. Our estimates of the stem ages of Nearctic scoliid clades are consistent with dispersal across Beringia during the Oligocene or later Eocene. Our study provides a foundation for future research into scoliid wasp evolution and biogeography by being the first to leverage genome-scale data and model-based methods. However, the precision of our dating analyses is constrained by the paucity of well-preserved fossils reliably attributable to the scoliid crown group. Despite concluding that the higher-level taxonomy of the Scoliidae is in dire need of revision, we recommend that taxonomic changes be predicated on datasets that extend the geographic and taxonomic sampling of the current study.

## Introduction

Members of the family Scoliidae, sometimes referred to as mammoth wasps, are large fossorial aculeates that comprise one of the most visually striking and easily identifiable hymenopteran clades. The family has a cosmopolitan distribution and includes approximately 560 described species (Osten, 2005). Adult mammoth wasps feed primarily on nectar, with honeydew (Illingworth, 1921) and possibly pollen (Jervis, 1998) also reported as food sources. The larvae develop as ectoparasitoids on the larvae of scarabaeid beetles (Clausen, 1940). Some studies have highlighted interesting aspects of mammoth wasp natural history, such as parasitism of ant inquilines (Burmeister, 1854; Jonkman 1980), pseudocopulation with orchids (Jones & Gray, 1974; Ciotek *et al*., 2006), fidelity of males to patrolling sites (Tani & Ueno, 2013), and efficient location of subterranean hosts (Inoue & Endo, 2008). Despite this, no study has attempted to reconstruct a phylogeny of the family, which precludes the examination of scoliid biology in an evolutionary context.

The lack of a solid phylogenetic hypothesis has also contributed to a lack of taxonomic clarity and stability. Day *et al*. (1981) referred to the group as “over-burdened nomenclatorially ”. Subsequently Argaman (1996), while describing the state of scoliid taxonomy as “disastrous ”, established a new subfamily, 21 new tribes, and 62 new genera without conducting a phylogenetic analysis. In assembling a checklist of all scoliid species, Osten (2005) ignored the taxonomic changes implemented by Argaman and implicitly synonymized many of the new taxa by placing their type species in other groups (Elliott, 2011; Kimsey & Brothers, 2016). Currently, the need for a thorough taxonomic revision is recognized (Elliott, 2011).

A robust phylogeny is a prerequisite for studies of character evolution, diversification patterns over time, and biogeography, as well as for a natural taxonomy. In turn, the lack of a stable natural taxonomy hampers research by making species determination difficult and by impeding the communication and indexing of scientific information. In the case of mammoth wasps, this is especially apparent in the context of their use as agents for the biological control of scarabaeid pests (Illingworth, 1921; Wilson, 1960; DeBach, 1964). Misidentification of the control agent (for an example, see Elliott (2011) on research by the Queensland Bureau of Sugar Experiment Stations) precludes the repeatability of research and past biological control attempts and means that valuable information discovered in the process cannot easily be traced to the right organism (Rosen, 1986). This is particularly unfortunate, since a large portion of what is currently known about scoliid development, phenology, and host interaction was discovered while evaluating and using mammoth wasps for biological control (e.g. Illingworth, 1921; Miyagi, 1960). In the process of updating the BIOCAT database of introductions of biological control agents, Cock *et al*. (2016) listed Scoliidae among the groups requiring further taxonomic work.

In the present study, we aim to establish a solid foundation for research into mammoth wasp evolution and systematics. We use ultraconserved element (UCE) sequence data (Faircloth *et al*., 2012; 2015) to infer scoliid phylogenetic trees using concatenation and under the multispecies coalescent (Rannala & Yang, 2003; Degnan & Rosenberg, 2009). Additionally, we leverage existing fossil data to estimate a timeline of scoliid evolution. To better understand potential biases resulting from model misspecification, we perform data filtering experiments based on matched-pairs tests of symmetry (Jermiin *et al*., 2017; Naser-Khdour *et al*., 2019) and assessments of model adequacy using data-based posterior predictive checks (Bollback, 2002; Huelsenbeck *et al*., 2001; Doyle *et al.,* 2015).

## Methods

### Taxon and locus selection

We successfully sequenced 85 specimens of Scoliidae for this study. Taxon selection was aimed at maximizing taxonomic and biogeographic diversity within the limits imposed by the availability of material from which DNA could be extracted. All biogeographic realms are represented, but with weaker sampling in Australasia and the Neotropical and Palearctic regions. We also included previously published data (Johnson *et al*., 2013; Faircloth *et al*., 2015; Branstetter *et al*., 2017a; Peters *et al*., 2018) from six additional scoliid specimens. See Table S1 for specimen collection data, voucher information, and resources used for taxonomic determination.

Based on an examination of morphology, we suspected that *Scolia bicincta* may constitute two separate species. We therefore sequenced multiple individuals from each putative species. However, given the focus of the current study on reconstructing the scoliid phylogeny and identifying major clades rather than on species delimitation, we retained only two specimens following a preliminary phylogenetic analysis (see below).

We used the bradynobaenid genus *Apterogyna* as the only outgroup, and mined UCE sequences (see “Sequence quality control, assembly, and UCE identification ” section below) from the partial genome published by Johnson *et al*. (2013). No sequences from other bradynobaenid taxa are publicly available, and we were unsuccessful in sequencing the specimens of *Bradynobaenus chubutinus* to which we had access. Bradynobaenidae is well-supported as the sister group to Scoliidae (Johnson *et al*., 2013; Branstetter *et al*., 2017a; Peters *et al*., 2018). It is a species-poor clade, making it easier to avoid highly disproportionate taxon sampling, which would be difficult if ants or apoids were used. Adding more distant outgroups also increases the chance that heterogeneity in the evolutionary process across lineages results in more severe violations of homogeneous phylogenetic models.

We used the hymenoptera-v2 ant-specific probe set (Branstetter *et al*., 2017b) targeting 2524 UCEs and 12 nuclear genes (“legacy ” markers).

### Wet lab methods

We extracted DNA from pinned and ethanol-preserved specimens using QUIAGEN DNeasy Blood & Tissue Kits. Extractions were semi-nondestructive. In the case of pinned specimens, we first removed them from their pins. For most specimens, we made holes in the right side of the thorax using an insect pin, then soaked the specimen in lysis buffer overnight. We used the buffer, now containing DNA, for subsequent extraction steps. We then washed the specimens in 95% ethanol and either dried and remounted them or returned them to ethanol. For especially large specimens (e.g. of *Megascolia*) we only used a sample of thoracic muscle for extraction. For some medium-to-large specimens that are part of longer collection series, we separated the metasoma and the head from the mesosoma, and soaked the mesosoma in lysis buffer overnight. In some cases, quantities of extraction reagents used had to be proportionally adjusted to accommodate specimen size. Finally, we either reassembled the specimen for remounting, or mounted the parts on separate points on the same pin.

We prepared, enriched, and pooled libraries using the hymenoptera-v2 ant-specific probe set following the protocols of Faircloth *et al*. (2015) as modified for use at the Ward Ant Lab (Ward & Branstetter, 2017). This was done in two separate batches. High-throughput sequencing was performed at the Huntsman Cancer Institute, University of Utah on an Illumina HiSeq 2500 platform (125 cycle paired-end) for the first batch and at the Novogene facility in Sacramento, CA on an Illumina HiSeq 4000 for the second batch.

### Sequence quality control, assembly, and UCE identification

After receiving demultiplexed reads, we used three different bioinformatics pipelines for quality control and *de novo* assembly.

#### Pipeline A

We performed quality-aware 3’ adapter trimming with Scythe (https://github.com/vsbuffalo/scythe) version 0.991. This was followed with 5’ adapter trimming with cutadapt (Martin, 2011) version 1.14 using a minimum overlap of 3 and an error tolerance of 0.16. We subsequently trimmed the reads with sickle (Joshi & Fass, 2011) version 1.33 using a quality threshold of 34 and a length threshold of 50. Assembly was done with Trinity (Grabherr *et al*., 2011) version 2.6.6 using a kmer size of 31. We also generated alternative assemblies with Velvet (Zerbino & Birney, 2008) version 1.2.10 and VelvetOptimiser (https://github.com/tseemann/VelvetOptimiser) version 2.2.4. However, the Velvet assemblies yielded significantly fewer UCE-containing contigs (data not shown, available upon request) and were not used for subsequent steps.

#### Pipeline B

We used HTStream (https://github.com/s4hts/HTStream) version 1.1.0 for adapter and quality trimming. The HTStream pipeline consisted of the following steps: (1) calculating basic statistics on the raw reads with hts_Stats (2) screening for phiX with hts_SeqScreener, (3) removing polyA/T sequences with hts_PolyATTrim with minimum size set to 100, (4) screening for adapter contamination with hts_SeqScreener using the i5 and i7 adapter sequences corresponding to each sample, with the kmer size set to 15, and with the percentage-hits argument set to 0.01, (5) a second round of adapter screening with hts_AdapterTrimmer, (6) quality-based 5’ and 3’ trimming with hts_QWindowTrim, (7) extracting the longest subsequences without “N ”s using hts_NTrimmer with the minimum length set to 50, and finally (8) calculating statistics on the processed reads with hts_Stats. In order to speed up read processing, we wrote a python script that can run the pipeline in parallel on more than one sample if the number of available CPU cores is at least twice the number of steps in the pipeline.

We then assembled the reads with Spades (Bankevich *et al*., 2012) using a wrapper script from the phyluce package (Faircloth, 2016), version 1.6.8. Except for increasing allowed memory usage, settings were left at phyluce defaults.

#### Pipeline C

We used Illumiprocessor (Faircloth, 2013), a wrapper around Trimmomatic (Bolger *et al*., 2014) and part of the phyluce package, for adapter and quality trimming. Spades was used for *de novo* assembly as in Pipeline B above.

For all pipelines, we used FastQC (Andrews, 2010) to evaluate reads before and after quality-control procedures.

We put reads from the first sequencing batch through Pipeline A and subsequently Pipeline B, while reads from the second sequencing batch were processed with Pipeline B and (with the exception of two samples) Pipeline C. In the case of ingroup taxa with previously published data (*Colpa sexmaculata*, *Colpa alcione*, *Proscolia* sp. EX568, *Scolia hirta*, *Scolia verticalis*, and Scoliinae sp. EX577), we used the available assemblies and did not redo quality control and assembly. In all cases, we used the phyluce_assembly_match_contigs_to_probes, phyluce_assembly_get_match_counts, phyluce_assembly_get_fastas_from_match_counts, and phyluce_assembly_explode_get_fastas_file scripts to identify UCE-containing contigs and write them to fasta files for downstream analyses.

Pipelines A and B recovered similar numbers of UCEs per sample, although Pipeline B resulted in assemblies with higher N50 as calculated in QUAST (Gurevich *et al*., 2013) version 5.0.2 on both whole assemblies and assemblies filtered to UCE-containing contigs only. Pipelines B and C were close in terms of both number of recovered UCEs and N50. See Tables S2-3 for details. However, each pipeline recovered some UCEs that the other pipelines did not. Therefore, we combined the assemblies, choosing the longer contig in cases where a contig containing the same UCE was recovered in both assemblies. However, longer contigs may either represent genuine sequence or be the result of assembly errors. We visually inspected alignments prior to most downstream analyses to identify and remove misaligned sequences possibly originating from misassembly.

Due to low UCE yield from some samples in the second sequencing batch (likely due to failed enrichment) and concerns over contamination, we did the following to identify problematic samples: (1) selected loci that were represented by > 75% of taxa, (2) aligned sequences from those loci using MAFFT (Katoh & Standley, 2013), (3) edge-trimmed the alignments using the phyluce_align_get_trimmed_alignments_from_untrimmed script from phyluce, and (4) estimated a phylogeny (Fig. S1) using maximum likelihood (ML) with IQTREE (Minh *et al*., 2020; Hoang *et al*., 2018; Chernomor *et al*., 2016; Nguyen *et al*. 2015) version 2.0-rc2 while partitioning by locus and filtering out loci using a matched-pairs test of symmetry (Jermiin *et al*., 2017; Naser-Khdour *et al*., 2019). Thirteen taxa associated with suspected failed enrichments clustered together in two “clades ” with very long branches, corroborating the spurious nature of the obtained sequences (Fig. S1). These taxa were not used in subsequent phylogenetic analyses and are not included in the counts under the taxon and locus selection section above.

In the case of *Apterogyna*, we mined UCE sequences from the partial genome of Johnson *et al*. (2013). We aligned UCE probes to the contigs using the phyluce_probe_run_multiple_lastzs_sqlite script from the phyluce package. We then extracted matching sequences in fasta format using the phyluce_probe_slice_sequence_from_genomes script, setting the flanking length to 700 bases.

### Phylogenetic analysis

Unless otherwise indicated, we performed all multiple-sequence alignments using MAFFT version 7.407 with the E-INS-i algorithm (Altschul, 1998). Preliminary visual inspection of alignments confirmed that they often contain multiple conserved, well-aligned regions separated by ambiguously aligned regions. This better conforms to the assumptions behind the E-INS-i algorithm. L-INS-i (Gotoh, 1993), on the other hand, assumes a single, contiguous alignable region. All edge-trimming was done using the phyluce_align_get_trimmed_alignments_from_untrimmed script. See log files (in repository listed under the data availability section below) for parameters used. All Bayesian phylogenetic analyses were performed using RevBayes (Höhna *et al*., 2014; 2016) version 1.0.12 unless otherwise indicated.

#### Analysis 1a

We performed a preliminary run combining all (non-spurious) data from both sequencing batches, including all *Scolia bicincta* samples. This helped inform which *S. bicincta* samples to retain, as discussed below. We selected loci that had no more that 20% missing data at the site level (after including taxa without data) and estimated a phylogeny using ML with IQTREE while partitioning by locus and filtering out loci using a matched-pairs test of symmetry (0.05 p-value cutoff).

#### Analysis 1b

We performed a second ML analysis with the goal of leveraging data from as many loci and taxa as possible while maintaining acceptable total levels of missing data. Given that analysis 1a indicated that samples of *S. bicincta* fall into two distinct clades that are sister to *S. dubia* and *S. mexicana* respectively (Fig. 1), we removed all but two *S. bicincta* samples (one from each putative species). In addition to phylogenetic position, the decision on which samples to retain was based on the number of recovered UCEs and on assembly quality statistics calculated using QUAST. We also removed *Scolia hirta* and Scoliinae sp. EX577, both from previously published studies, because they had very high fractions of missing data. After taxon removal, we redid alignment and edge-trimming. We then sorted loci by increasing fraction of missing data at the site level and progressively selected loci until the cumulative fraction of missing data reached 25% (1235 loci were selected at this point). After filtering using tests of symmetry in IQTREE, we retained 727 loci. We concatenated the alignments and selected a substitution and across-site rate variation (ASRV) model (from a pool of substitution models from the GTR (Tavaré, 1986) family and discretized gamma (Yang, 1994) and free-rates ASRV models) for each locus based on Bayesian Information Criterion (BIC) (Schwarz, 1978) scores. We then estimated a phylogeny and performed 1000 ultrafast bootstrap replicates while leaving other IQTEE settings at default.

**Figure 1.**
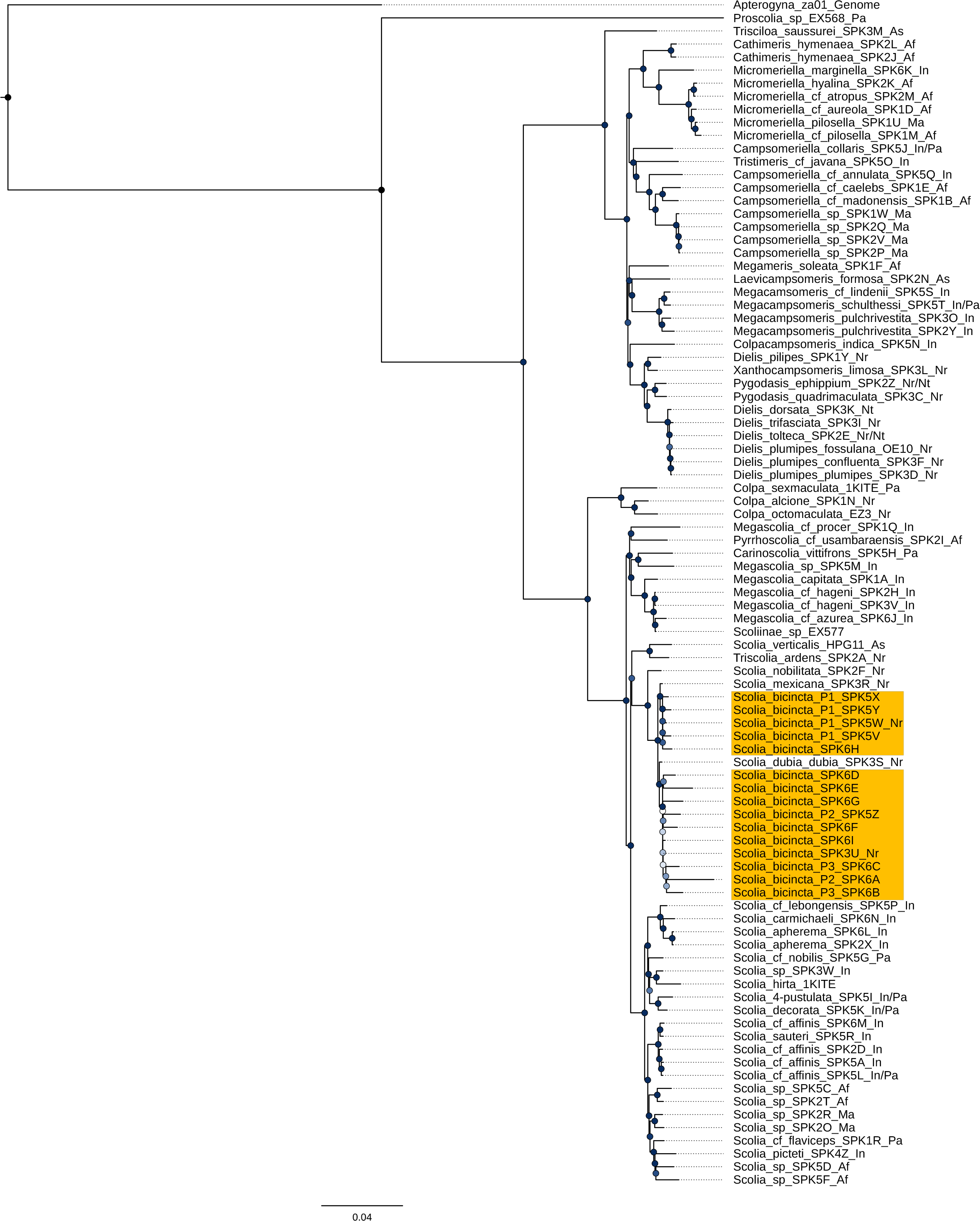
Maximum likelihood phylogeny including all samples (analysis 1a). Two distinct clades of *Scolia bicincta* are highlighted in yellow. Node support values are based on Ultrafast Bootstrapping in IQTREE; darker nodes reflect higher support.

#### Analysis 1c

In order to account for potential gene-tree-gene-tree conflict due to incomplete lineage sorting, we estimated species trees using the program ASTRAL-MP version 1.15.1 (Yin *et al.,* 2019). Starting with the same set of taxa used in analysis 1b, we redid alignment and edge-trimming, discarding alignments shorter than 600 bases. Given that highly fragmentary sequences can negatively affect accuracy (Sayyari *et al*., 2017), we subsequently removed taxa with more than 50% missing data and discarded alignments that retained fewer than 66 taxa. We then inferred gene trees using IQTREE with model selection settings similar to those in analysis 1b above while also performing matched-pairs tests of symmetry. We based subsequent species tree inference on three sets of gene trees: The first set contained trees corresponding to all loci, the second contained only trees from loci that failed the maximum test of symmetry (p-value < 0.05), and the last contained only trees from loci that passed.

Additionally, we estimated posterior distributions of gene trees in a Bayesian framework under the GTR+G model, followed by posterior predictive simulation (Bollback, 2002; Brown, 2014b; Doyle *et al.,* 2015; Höhna *et al.,* 2018) and calculation of posterior predictive p-values using two test statistics: multinomial likelihood (Goldman, 1993; Bollback, 2002) and chi-squared (Huelsenbeck *et al*., 2001; Foster 2004). Similarly to the ML-based analyses above, we then used different sets of maximum clade credibility (MCC) gene trees and gene tree posterior distribution samples (3000 trees per gene) for species tree inference with ASTRAL-MP. Using an alpha of 0.05 and the Bonferroni correction to account for multiple testing, we treated loci for which the posterior predictive p-value with either test statistic was < 0.025 as loci for which the model was likely inadequate. This set included 922 of the total 954 loci. For each test statistic, we also split the loci into two sets, each respectively representing loci with the lowest and highest effect sizes for that statistic. Finally, we created another similar pair of sets but based on the Pythagorean sum of the effect sizes for both statistics. When using gene tree posterior distribution samples with ASTRAL, we performed bootstrapping using the -b option and set the number of replicates to 1000.

In datasets used for analyses 1a-c, *Apterogyna* and *Proscolia* have disproportionately high fractions of missing data (49% and 73% respectively) compared to other taxa. However, removing these taxa means the loss of the only outgroup. We therefore took a two-step approach: First, we performed an analysis (2a) only using loci with data available from both *Apterogyna* and *Proscolia* to minimize the potential impact of missing data on the inferred position of the root as well as on the placement of *Proscolia*. However, significantly cutting down the base dataset could result in loss of resolution in some parts of the tree. To address this, we performed another set of analyses (analyses 3a and 3b; see below) excluding *Apterogyna* and *Proscolia* as well as loci used in analysis 2a but conditioning on the position of the root inferred in analysis 2a. This allowed use of the remaining majority of the original data to resolve relationships within Scoliidae.

#### Analysis 2a

We started with the same taxon set as for analysis 1b and selected aligned, trimmed fasta files corresponding to the 647 loci that have sequences from both *Apterogyna* and *Proscolia*. We used the biclustering algorithm of Uitert *et al*. (2008) as implemented in the R (Core R Team, 2020) package BicBin (https://github.com/TylerBackman/BicBin) to find large, dense biclusters of taxa and loci. We chose a set of 68 taxa and 484 loci with >99% completeness (presence or absence of sequence for a given taxon and locus pair treated as a binary value). We then retrieved unaligned, untrimmed fasta files corresponding to the above loci and removed the taxa that are not part of the selected set. The sequences were then aligned and edge-trimmed. Given that the phylogenetic models we planned to use do not directly model indels (gaps are treated as missing data) and that unique indels are unlikely to contribute significant information, we removed all unique indels (i.e. columns where all taxa except one are represented by a gap) from the alignments. Calculating basic alignment statistics using AMAS (Borowiec, 2016) and visually inspecting the alignments in AliView (Larsson, 2014) revealed that *Apterogyna* sequences were (1) sometimes much shorter than those of other taxa for a given locus and (2) sometimes had poorly aligned sections. We therefore only retained alignments containing at least 500 non-ambiguous bases for both *Apterogyna* and *Proscolia*. We then manually trimmed alignment edges that contained no *Apterogyna* sequence and also trimmed any parts with suspected alignment uncertainty while discarding alignments that were poor throughout their length. Any alignments that became shorter than 300 bases were also discarded.

In order to assess model adequacy on the remaining 177 loci, we performed Bayesian phylogenetic analyses under the GTR+G model followed by posterior predictive simulation on each locus individually using the program RevBayes. We calculated the multinomial likelihood and chi-squared (as applied to nucleotide composition across taxa) test statistics and associated posterior predictive p-values and effect sizes on the empirical and simulated data using custom R code. For the purpose of filtering data for which the available model is suspected of being inadequate, one must choose some threshold. In advance of looking at the output, we decided to use an overall alpha of 0.05 and use the Bonferroni correction to account for multiple testing. We therefore discarded loci for which the posterior predictive p-value with either test statistic was < 0.025. We concatenated the remaining 31 alignments and used them for phylogeny estimation. Each locus was assigned a separate GTR+G substitution model and tree length parameter (i.e. branch length multiplier), while a single vector of branch lengths drawn from a flat Dirichlet prior was shared among partitions. See used Rev scripts for further details. We assessed convergence for numerical parameters through visualization of posterior samples in Tracer (Rambaut *et al*., 2018) version 1.7. For tree topologies, we made plots comparing posterior probabilities of splits across both runs using the bonsai (May & Moore, 2017) version 0.9 R package and calculated the Average Standard Deviation of Split Frequencies (ASDSF).

#### Analysis 2b

Rasnitsyn (1993) identified only one fossil from Shangwang, Shandong, China as unequivocally belonging to the scoliid crown group. This fossil was attributed by Zhang (1989) to the extant species *Scolia prismatica*, currently in the genus *Megacampsomeris*. Yu *et al*. (2021) dated the Shanwang shale to approximately 18.5 Ma, in the early Miocene. Species described in later studies (Rasnitsyn & Martınez-Delclos, 1999; Nel *et al*., 2013; Zhang *et al*., 2015) are either connected to the crown Scoliidae by venation characters alone, or are of uncertain placement. This limits the information available to precisely estimate divergence times. Given this limitation and our inability to examine the *M. prismatica* specimen, we chose a conservative approach and estimated a broad timeline of scoliid evolution by calibrating the node representing the most recent common ancestor of Scoliidae and Bradynobaenidae using the age of *Protoscolia normalis*, a putative stem scoliid dated to approximately 125.5 Ma (Haichun *et al*., 2002). We started with 177 processed alignments from analysis 2a (i.e. the state of the dataset after removal of short alignments but prior to filtering using posterior predictive checks). We then performed analyses on individual loci followed by posterior predictive simulation. We used a birth-death prior on tree topologies and node ages with a scaled beta prior on the root age (125.5 Ma minimum age, 174.1 Ma maximum age, 132.5 Ma expected age, and a standard deviation of 5.5 Ma) and an uncorrelated lognormal relaxed clock model. See used Rev scripts for further details. After filtering loci in a similar manner to what was done in analysis 2a, we concatenated and analyzed the remaining 63 loci, adding rate multiplier parameters to allow the overall substitution rates to vary among loci.

In addition to the conservative primary analysis, we tested the effect of calibrating additional nodes using fossils of less certain placement. Although it is doubtful that the fossil described by Zhang (1989) belongs to an extant species, for the first additional analysis, we used it to set an 18.5 Ma minimum age (lognormal node age “prior ” offset by 18.5, with a mean of 5.0 (mu ≈ 1.44) relative to the offset and a sigma of 0.587405) for the *Megacampsomeris* clade. For the second analysis, we used both the *Megacampsomeris* calibration above as well as a calibration of the scoliid crown group age based on *Araripescolia magnifica* (Nel *et al*., 2013) (lognormal node age “prior ” offset by 112.6 Ma, a mean of 10.0 relative to the offset and a sigma of 0.587405).

#### Analysis 2c

In order to account for potential gene-tree-gene-tree conflict due to incomplete lineage sorting, we performed a species tree estimation analysis under the multispecies coalescent (e.g. Rannala & Yang, 2003; Degnan & Rosenberg, 2009) using the BEAST2 (Bouckaert *et al*., 2019) package STACEY (Jones, 2017). We used the same 63 loci from analysis 2b. Collapse weight was drawn from a beta prior with an alpha of 1.0 and a beta of 19.0 (mean 0.05, to reflect the belief that most samples are likely from distinct species). We used a lognormal prior on the popPriorScale parameter with a mean and standard deviation (in real space) of 1.0E-6 and 2.0 respectively. We enabled estimation of the relative death rate, which in this context corresponds to using a birth-death (as opposed to Yule) tree prior, and used a strict clock model. The site model was set to GTR+G, unlinked among loci. We ran four independent chains and combined and summarized the output using the logcombiner and treeannotator tools packaged with BEAST2.

#### Analysis 2d

We additionally performed species tree estimation using ASTRAL-MP. We used the same 177 starting loci from analysis 2a, but reran Bayesian gene tree estimation and posterior predictive simulation after removing taxa which had no data for a given locus. We then assembled sets of loci based on posterior predictive effect sizes in a manner similar to that in analysis 1c.

#### Analysis 3a

In order to leverage more data to resolve relationships within Scoliidae, we set up an analysis that conditions on the position of the root inferred in analysis 1b while removing *Proscolia* and *Apterogyna* from the dataset. We followed a locus and taxon selection, alignment, and trimming procedure similar to that in analysis 2a. We chose a set of 72 taxa and 617 loci at 91% completeness from a pool of loci that excludes those used in analysis 1b. After discarding all alignments that, after trimming, were shorter than 300 bases or had more than 25% missing data at the site level, 469 alignments were retained. We did not trim alignments manually at this stage as the number of loci was large and the exclusion of *Apterogyna* and *Proscolia* improved alignment quality (assessed by visual inspection of a subset of alignments). We then ran Bayesian phylogenetic analyses followed by posterior predictive simulation on each individual alignment as in 2a. All alignments which passed filtering, as well as some that did not, were visually evaluated, and in a few cases problematic regions were manually trimmed. One locus was excluded due to very poor alignment. We reran posterior predictive tests on all alignments that have been altered. We then performed a concatenated analysis analogous to that in 2a, which included all loci that passed filtering and were not subsequently edited and loci which were edited and subsequently passed filtering.

#### Analysis 3b

The data processing and phylogenetic analysis procedures were analogous to those of analysis 3a, except we used a birth-death prior on trees and node ages (with no node calibration and with the root age arbitrarily fixed to 100 units) and an uncorrelated lognormal clock model.

## Results

### Sequence quality control, assembly, and UCE identification

Using pipeline A (Scythe + cutadapt + sickle + Trinity), we recovered 1941.9 UCE-containing contigs on average across specimens from batch 1, which is almost identical to the 1943.7 UCE-containing contigs recovered when using pipeline B (HTStream + Spades). However, the output of pipeline B had higher average N50 (2112.6 versus 1191.4, calculated from on-target contigs only) and a higher average number of UCE-containing contigs longer than 1000 bases (1429.0 versus 968.3).

The differences between outputs from pipelines B and C applied to specimens from batch 2 were in some ways less pronounced. The average number of UCE-containing contigs was 1726.3 and 1785.6 for pipelines B and C respectively, while average values for N50 were 1477.8 and 1482.5 respectively. The average number of on-target contigs longer than 1000 bases was 1036.0 for pipeline B and 1077.6 for pipeline C. When calculating these statistics, we excluded batch 2 samples for which we suspected failed enrichment (see corresponding section under Methods for details and SI tables 1 and 2 for full QUAST statistics).

Overall, we recovered a total of 2495 UCE loci and an average of 1883.6 UCE loci per taxon across 91 taxa (including 6 taxa from previously published studies).

### Phylogenetic analysis

#### Analysis 1a

A total of 176 loci were retained after all filtering steps and used to estimate a phylogeny by maximum likelihood (Fig. 1). We recovered *Proscolia* as the sister group to all remaining Scoliidae, which correspond to the subfamily Scoliinae *sensu* Day *et al*. (1981). The tribe Scoliini is monophyletic. However, in contrast to the assumptions behind the current scoliid taxonomy (Osten, 2005), the genus *Colpa* was recovered as sister to the Scoliini, rendering the Campsomerini paraphyletic.

A clade represented by the scoliine genera *Megascolia*, *Pyrrhoscolia*, and *Carinoscolia* is sister to all other Scoliini, which in turn form three distinct groups. All New World members of the genus *Scolia* form a clade. We recovered *Scolia verticalis*, an Australasian species, as sister to the morphologically unusual Nearctic species *Triscolia ardens*. Given the unexpected nature of this pairing, we conducted an additional analysis (see Supporting Information for details) using (1) the “legacy ” markers enriched from *T. ardens* as part of this study and from *S. verticalis* (from Faircloth *et al*. (2015), the source of *S. verticalis* UCE data used in this study), (2) corresponding Sanger data from the same specimen of *S. verticals* (Brady *et al*., 2006; Ward & Fisher, 2016), and (3) corresponding Sanger data from different specimens of *T. ardens* (Pilgrim *et al*., 2008) and *S. verticalis* (Klopfstein & Ronquist, 2013).

Sequences from the specimens used in this study grouped with their corresponding sequences from independent samples (Fig. S2), which makes contamination or data curation errors a less likely explanation for the relationship between *T. ardens* and *S. verticalis* inferred here. All remaining sampled Scoliini form an Old World clade that is sister to the clade consisting of New World *Scolia* + (*T. ardens* + *S. verticalis*).

Samples of *Scolia bicincta* fall into two separate clades: one sister to *Scolia mexicana* and the other sister to *Scolia dubia*. This suggests the two groups belong to different species.

Campsomerini minus *Colpa* (provisionally referred to as Campsomerini *sensu stricto* from here on) is monophyletic. *Trisciloa saussurei* (not to be confused with members of the genus *Triscolia*) is inferred to be the sister taxon to the remaining Campsomerini *sensu stricto*. Within the latter group, all sampled New World taxa form a single clade. The closest relative of this New World clade is the Indomalayan taxon *Colpacampsomeris indica*, followed by a clade including the Afrotropical *Megameris soleata*, the Australiasian *Laevicampsomeris formosa*, and the Indomalayan genus *Megacampsomeris*. *Megacampsomeris* itself is recovered as monophyletic. Taxa occurring in Madagascar, such as *Micromeriella pilosella* and some *Campsomeriella*, have their closest affinities with Afrotropical taxa but do not form a monophyletic group.

#### Analysis 1b

We used 727 loci from 76 taxa to reconstruct the tree in Fig. 2. The results are largely congruent with those from analysis 1a above, with the exception of the *Triscolia ardens* + *Scolia verticalis* group being recovered as sister to the Old World Scoliini (minus *Megascolia* + *Pyrrhoscolia* + *Carinoscolia*) as opposed to sister to the New World *Scolia*. *Colpa* is still recovered as sister to the Scoliini. The non-monophyly of *Dielis*, due to *Dielis pilipes* being more closely related to *Xanthocampsomeris* than to other *Dielis*, is likewise corroborated.

**Figure 2.**
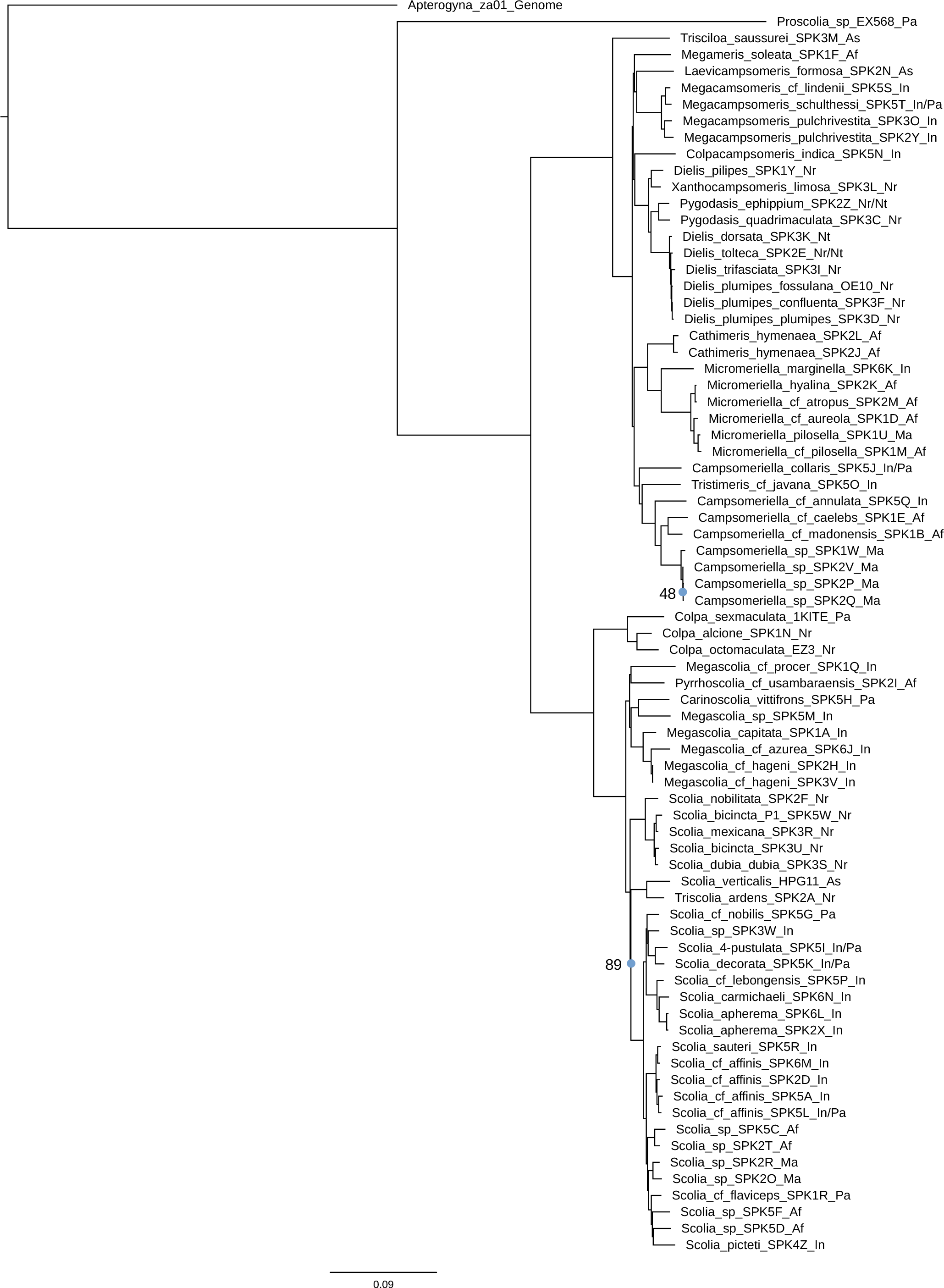
Maximum likelihood phylogeny based on 727 UCE loci (analysis 1b). Node support values based on Ultrafast Bootstrapping in IQTREE. All unlabeled internal nodes have 100% bootstrap support.

#### Analysis 1c

For all analyses, the topology of the “main ” ASTRAL tree (based only on ML or MCC gene trees) was effectively the same as the consensus topology estimated using gene tree posterior distributions and bootstrapping. Differences were limited to quadripartitions with very low support (e.g. 0.46 local posterior probability for most probable resolution, versus 0.35 for the next most probable alternative) or to relationships within species (e.g. *Dielis plumipes*).

The inferred topology based on ML trees from all loci (Fig. 3C) agrees with that from analysis 1b above. The topology based only on loci not failing the maximum test of symmetry (Fig. 4, Fig. 3A) is identical, but with reduced support for the quadripartition involving *Megameris soleata*, *Laevicampsomeris formosa* + *Megacampsomeris*, *Colpacampsomeris indica* + New World Campsomerini, and the remaining Campsomerini. The topology inferred from loci failing the symmetry test (Fig. 3B) maintained high support for this quadripartition. On the other hand, the position of *Triscolia ardens* + *Scolia verticalis* became more uncertain, with 0.50 local posterior probability for the same placement as the other analyses above and 0.30 local posterior probability for *Triscolia ardens* + *Scolia verticalis* being sister to the New World *Scolia*.

**Figure 3.**
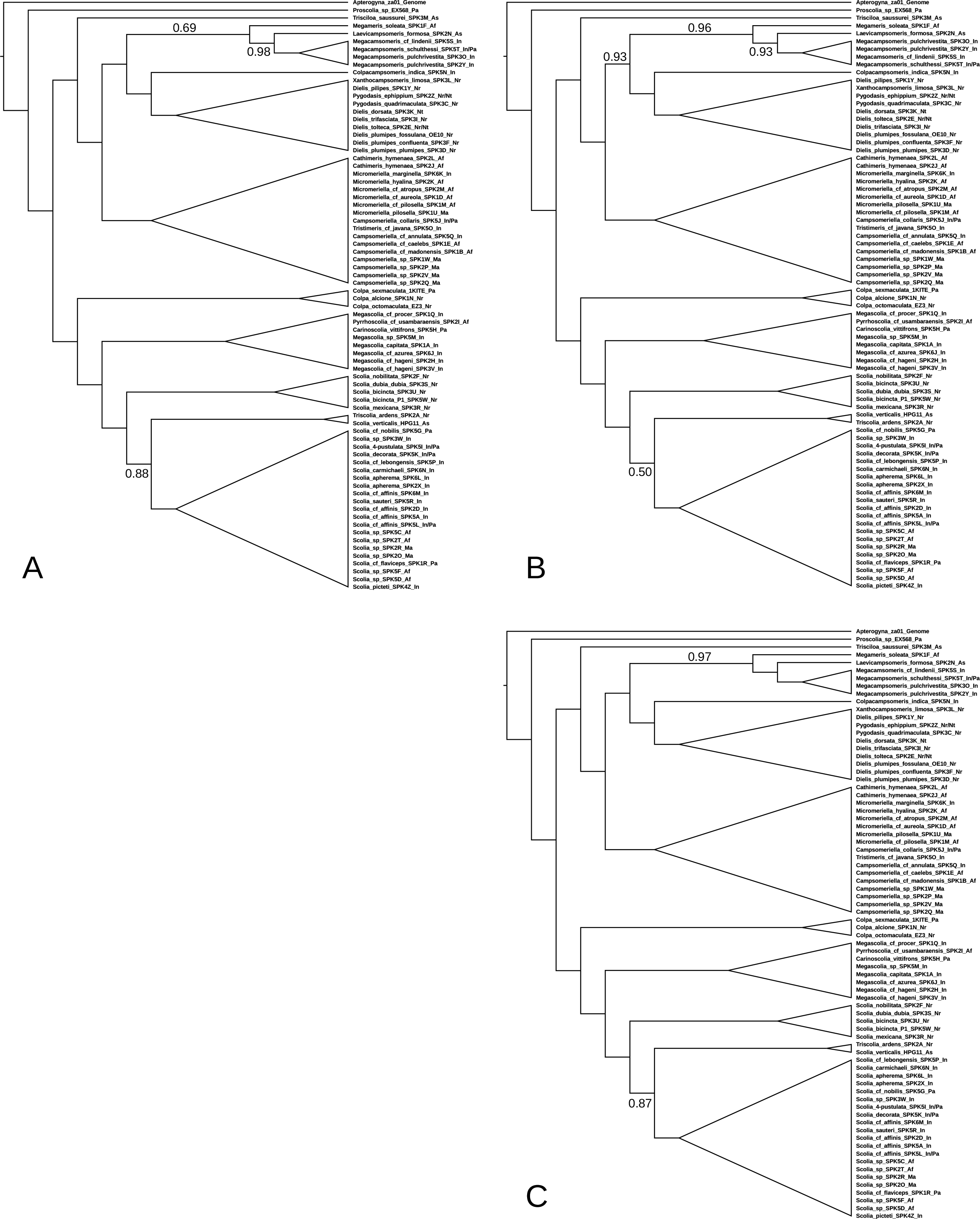
ASTRAL species trees (analysis 1c) based on ML trees of (A) loci not failing the maximum test of symmetry, (B) loci failing the the maximum test of symmetry, and (C) all loci. Branch labels represent the local posterior probability of the associated quadripartition. All unlabeled quadripartitions have a local posterior probability of 1.0.

**Figure 4.**
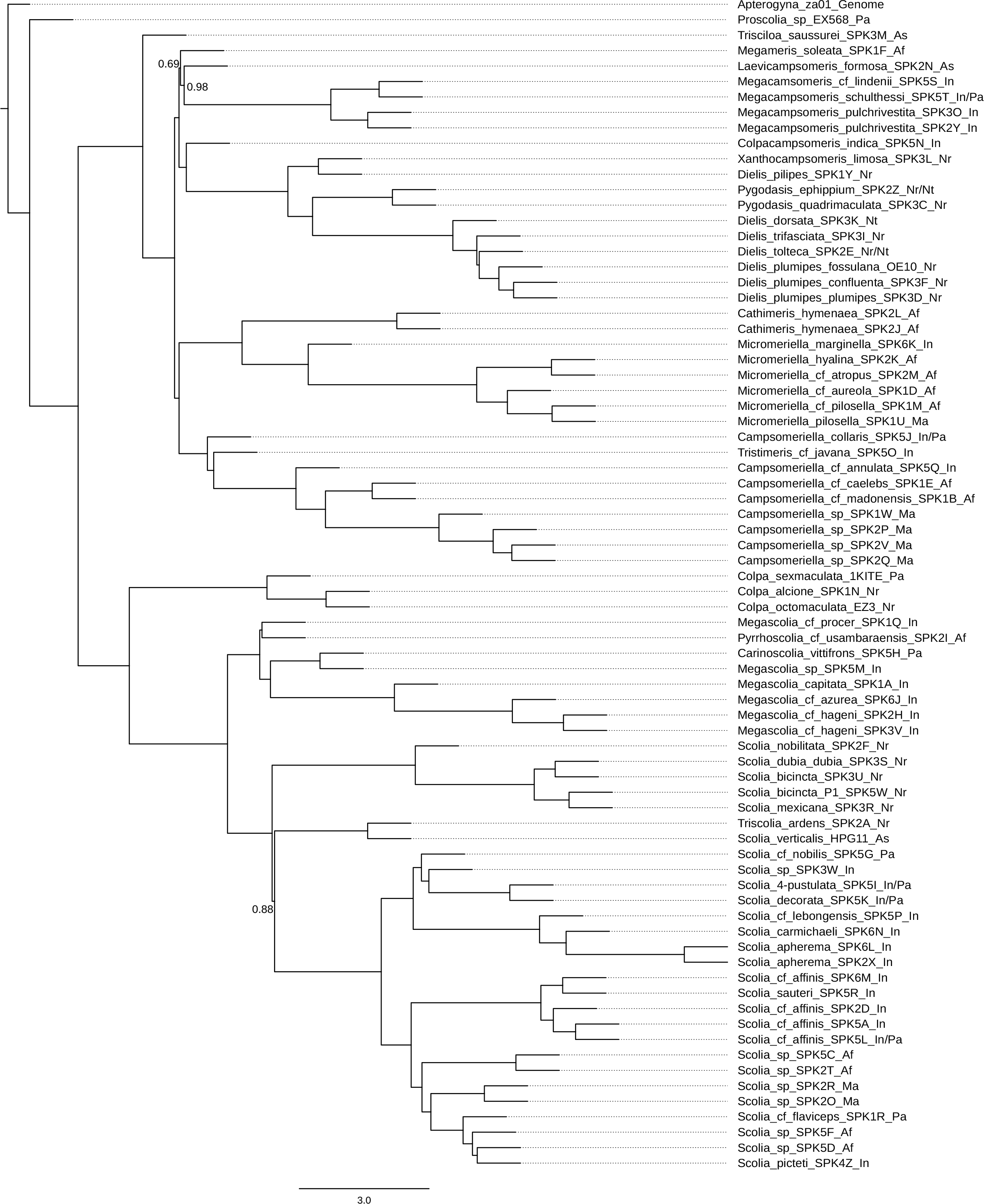
ASTRAL species tree (analysis 1c) based on ML trees of loci not failing the maximum test of symmetry. Branch labels represent the local posterior probability of the associated quadripartition. All unlabeled quadripartitions have a local posterior probability of 1.0.

Results of the analysis using MCC trees (as a way of summarizing tree posterior distributions) from all loci (Fig. 5D) agree with the ML-based results above with respect to the Campsomerini *sensu stricto*. However, the placement of *Triscolia ardens* + *Scolia verticalis* is not resolved, with 0.47 and 0.46 local posterior probability for a sister relationship with the sampled Old World *Scolia* and with the New World *Scolia* respectively. The ASTRAL tree based on loci with the lowest combined posterior predictive effect sizes (Fig. 5A) is similar to the tree above, with 0.46 local posterior probably in favor of (*Triscolia ardens* + *Scolia verticalis*) + New World *Scolia*, but a slightly lower probability (0.36) in favor *Triscolia ardens* + *Scolia verticalis* being sister to the Old World *Scolia*. The analysis of loci with highest combined posterior predictive effect sizes (Fig. 5B) resulted in stronger (0.82 local posterior probability) support for the (*Triscolia ardens* + *Scolia verticalis*) + New World *Scolia* hypothesis.

**Figure 5.**
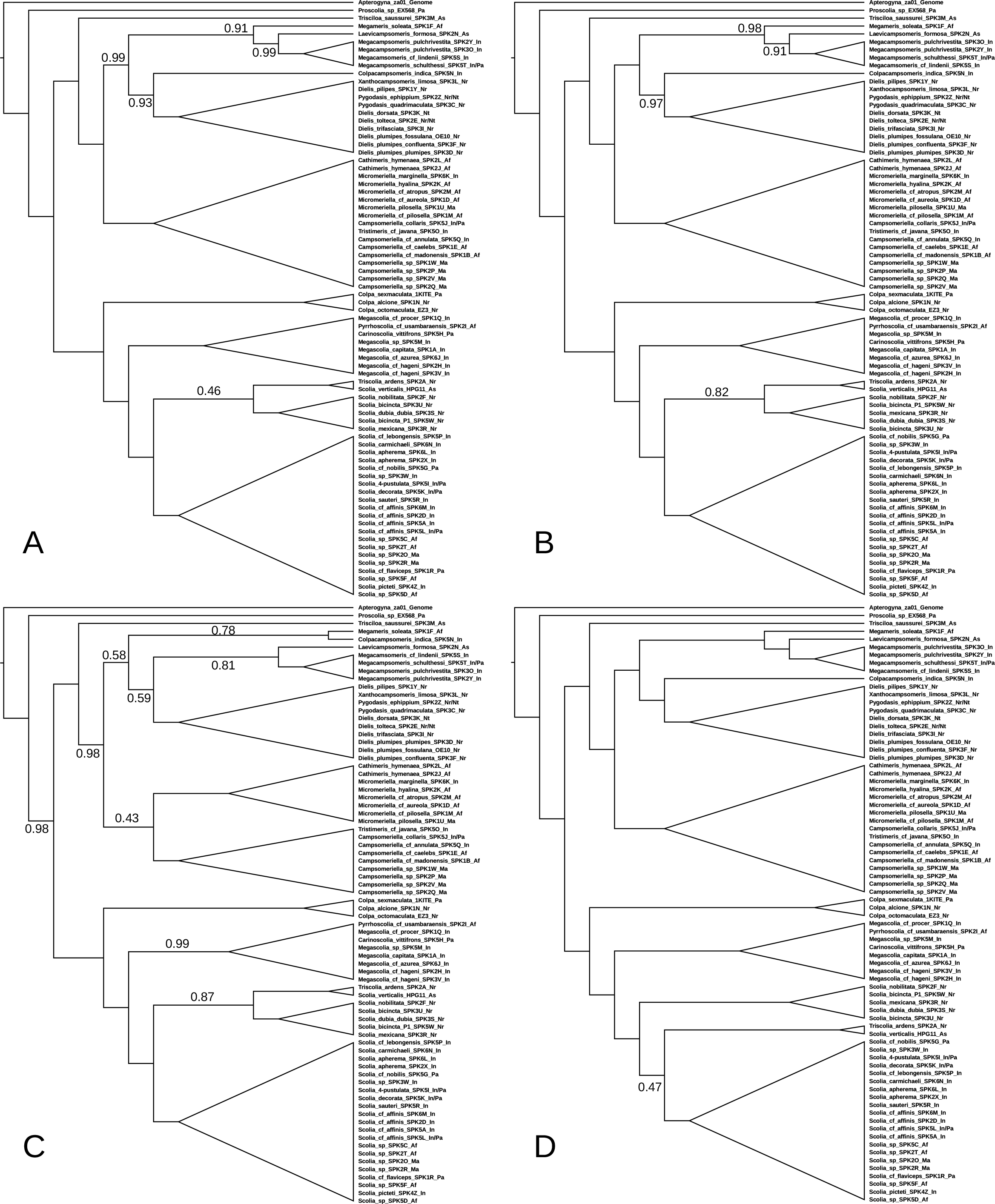
ASTRAL species trees (analysis 1c) based on MCC trees of (A) loci having the lowest (1/3) combined posterior predictive effect sizes, (B) loci having the highest (1/3) combined posterior predictive effect sizes, (C) loci for which the model was not found to be inadequate (alpha = 0.05), and (D) all loci. Branch labels represent the local posterior probability of the associated quadripartition. All unlabeled quadripartitions have a local posterior probability of 1.0.

Unexpectedly, this relationship was likewise supported (0.87 local posterior probability) when using only the 32 loci for which the model was not found to be inadequate (Fig. 5C) using posterior predictive checks, but resolution within the Campsomerini was significantly reduced. Crucially, all analyses agree with respect to the placement of *Colpa* as sister to the Scoliini.

#### Analysis 2a

The tree in Fig. 6 is the Maximum *A Posteriori* (MAP) tree summarized from two independent runs based on 31 loci for which the model was not found to be inadequate. The MCMC exhibited good convergence with respect to topology (see Fig. 7A for a comparison of split frequencies between runs). The average standard deviation of split frequencies was approximately 0.001.

**Figure 6.**
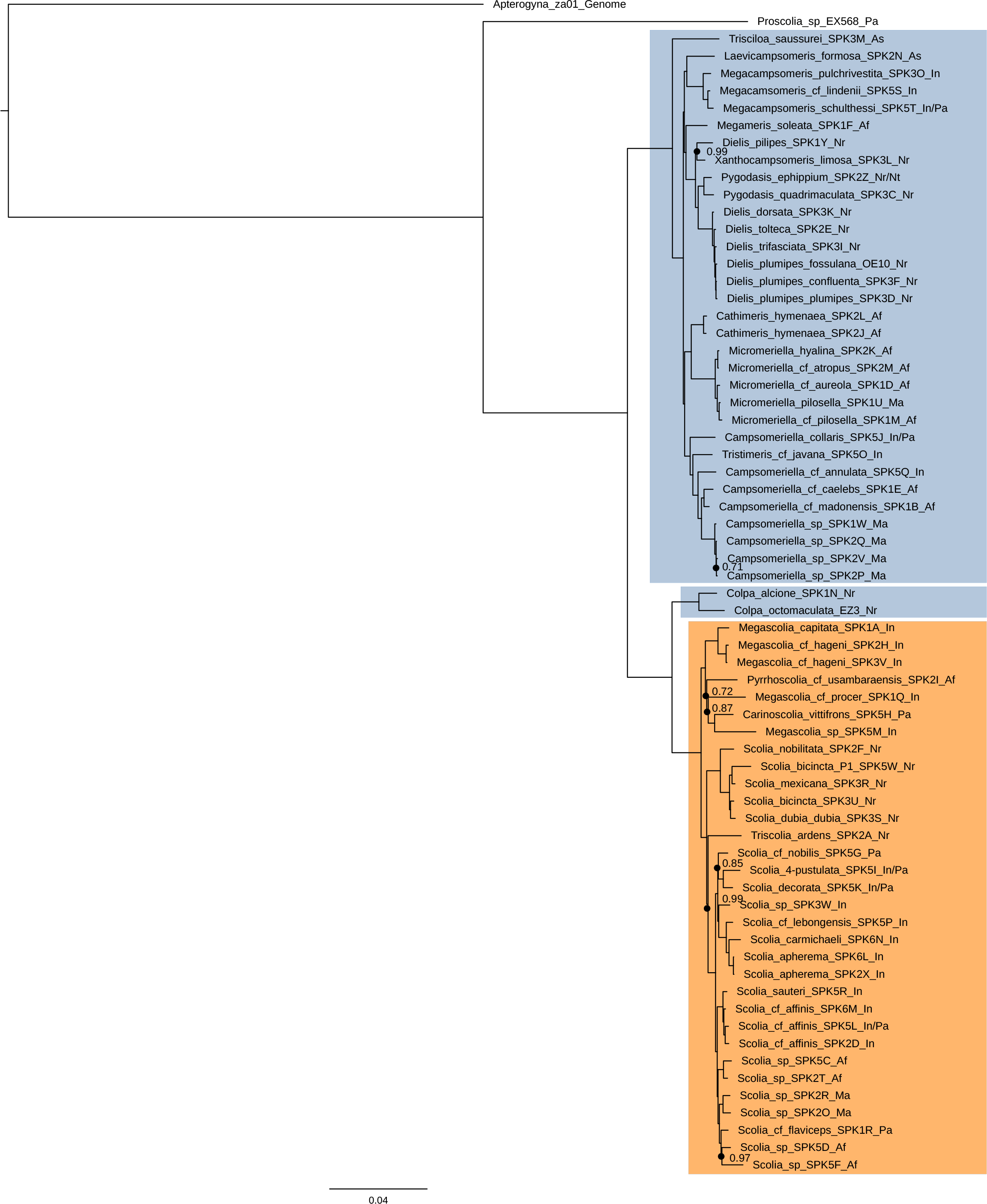
Bayesian MAP tree based on 31 UCE loci after data filtering using posterior predictive checks (analysis 2a). All unlabeled internal nodes have posterior probabilities of 1.0. Paraphyletic Campsomerini highlighted in blue; Scoliini highlighted in orange.

**Figure 7.**
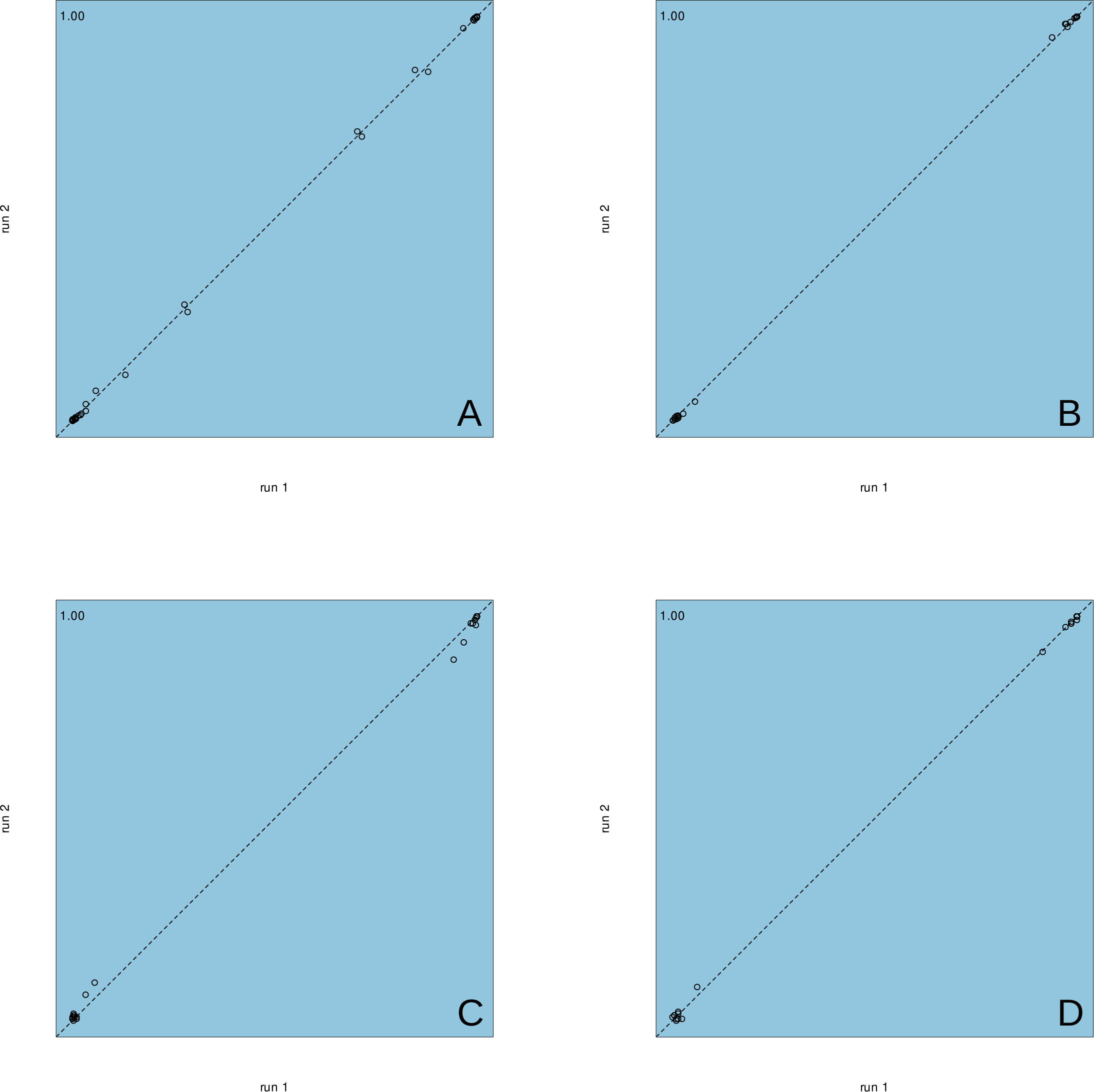
Comparison of split posterior probabilities between two independent MCMC runs: (A) analysis 2a; (B) analysis 2b, root calibration only; (C) analysis 2b, root + *Megacampsomeris* calibration; (D) analysis 2b, root + *Megacampsomeris* + Scoliidae calibration. Numbers in the top left corners represent R^2^.

This analysis places emphasis on reducing missing data in the outgroup and in *Proscolia*, removing poorly aligned sites, and reducing potential model violation at the expense of dataset size. Despite this, the tree backbone is fully resolved, with only a few shallow nodes having lower posterior probabilities. With respect to the position of the root, the results corroborate those from analyses 1a, 1b, and 1c:

*Proscolia* is sister to the Scoliinae, *Colpa* is sister to the Scoliini, and Campsomerini *sensu stricto* is sister to Scoliini + *Colpa*, with Campsomerini in the traditional sense being non-monophyletic. The position of *Triscolia* as sister to an Old World scoliine clade is congruent with that in analysis 1b but not analysis 1a. *Scolia verticalis*, which was recovered as sister to *Tricolia ardens* in previous analyses, was not represented here and in subsequent analyses due to a high proportion of missing data. While *Colpacampsomeris indica* was likewise excluded from this analysis for the same reason, *Megameris soleata* is placed as sister to the New World Campsomerini instead of being sister to *Laevicampsomeris* + *Megacampsomeris* as in analyses 1a and 1b.

#### Analysis 2b

A total of 63 loci were retained post-filtering and used to construct a chronogram (Fig. 8). See Fig. 7B for a plot of split frequencies from two independent runs. While most clade posterior probabilities are close to 1 and none are lower than 0.94, node age credible intervals are broad due to only one calibration point being available. The crown Scoliini are inferred to have likely originated after the Cretaceous-Paleogene (K-Pg) extinction event. The mean estimated crown ages of Campsomerini *sensu stricto* and of Scoliini + *Colpa* are 49 million years (Ma) and 58 Ma respectively, although the associated 95% highest posterior density (HPD) intervals extend past the K-Pg boundary. The mean estimated age of crown Scoliinae is 84 Ma, with lower and upper bounds of the 95% HPD interval at 56 Ma and 107 Ma respectively. The crown Scoliidae as a whole (and thus the split between Proscoliinae and Scoliinae) has a 95% HPD age interval bounded by 96 Ma and 145 Ma, placing the likely origin of the group in the Early Cretaceous.

**Figure 8.**
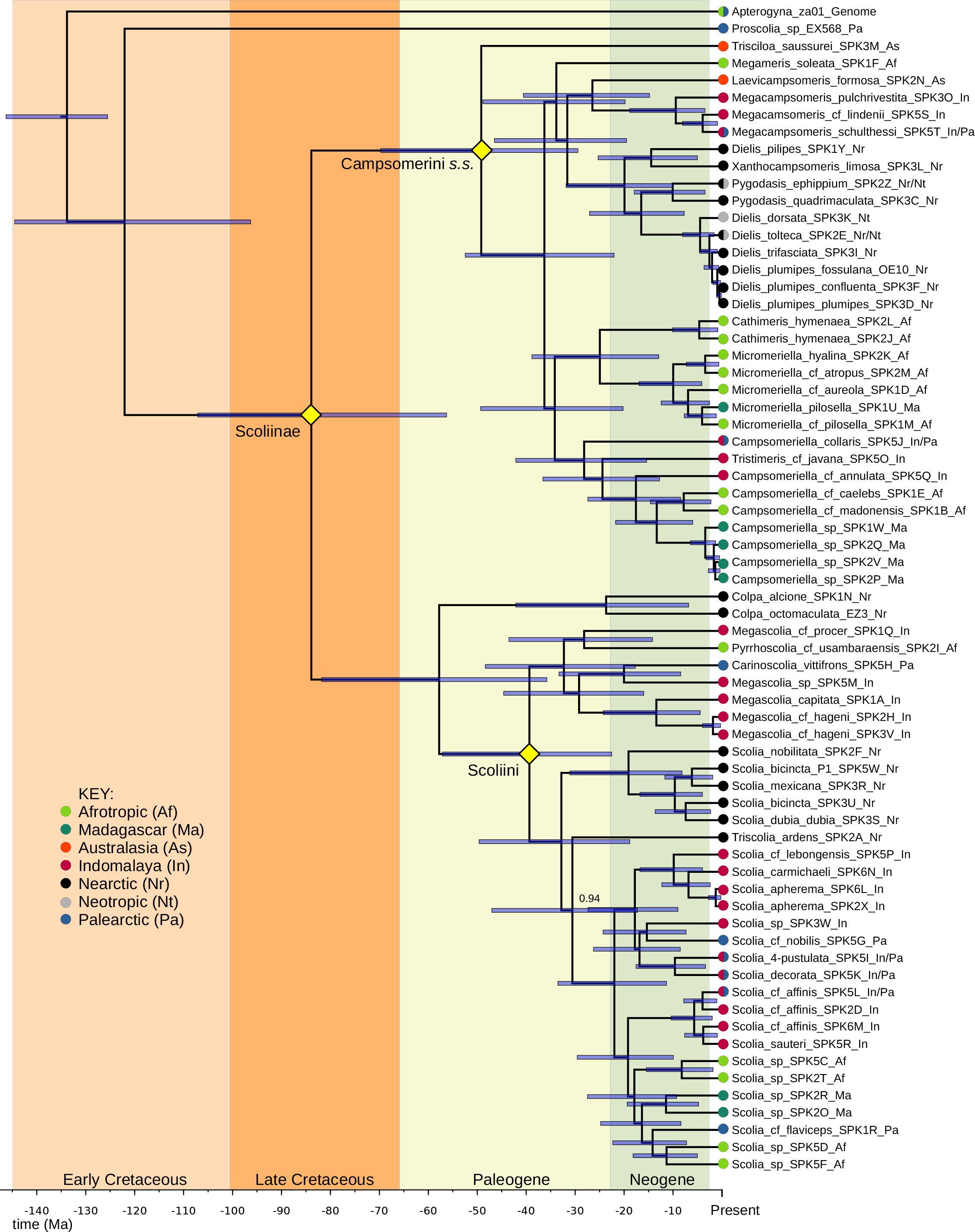
Bayesian MAP chronogram based on 63 loci after data filtering using posterior predictive checks (analysis 2b). Node bars represent age 95% HPD intervals. All unlabeled internal nodes have posterior probability of 0.97 or greater. Taxonomic labels indicated with 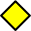.

Results from the analyses including additional fossil calibrations (Fig. 9-10) were broadly congruent with the results above, but with greater ages estimated for most nodes after the Scoliinae/Proscoliinae split. When using both additional calibrations, the posterior distributions of ages for Campsomerini *sensu stricto* and of Scoliini + *Colpa* had means of 63 Ma and 69 Ma respectively, with more posterior mass on pre-K-Pg ages compared to the more conservative analysis above.

**Figure 9.**
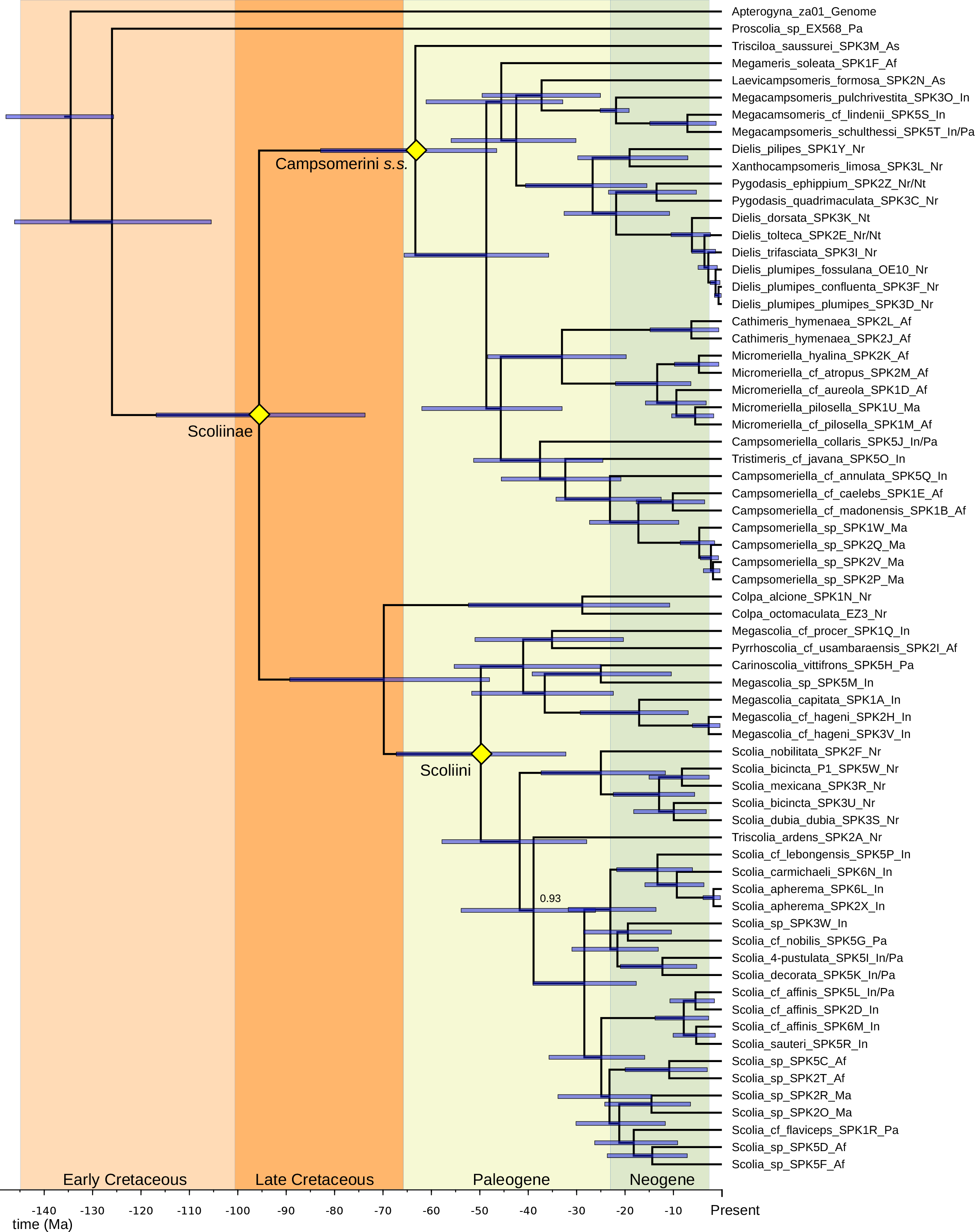
Bayesian MAP chronogram using additional calibration on crown *Megacampsomeris* (analysis 2b). All unlabeled internal nodes have posterior probability of 0.96 or greater. Taxonomic labels indicated with 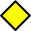.

**Figure 10.**
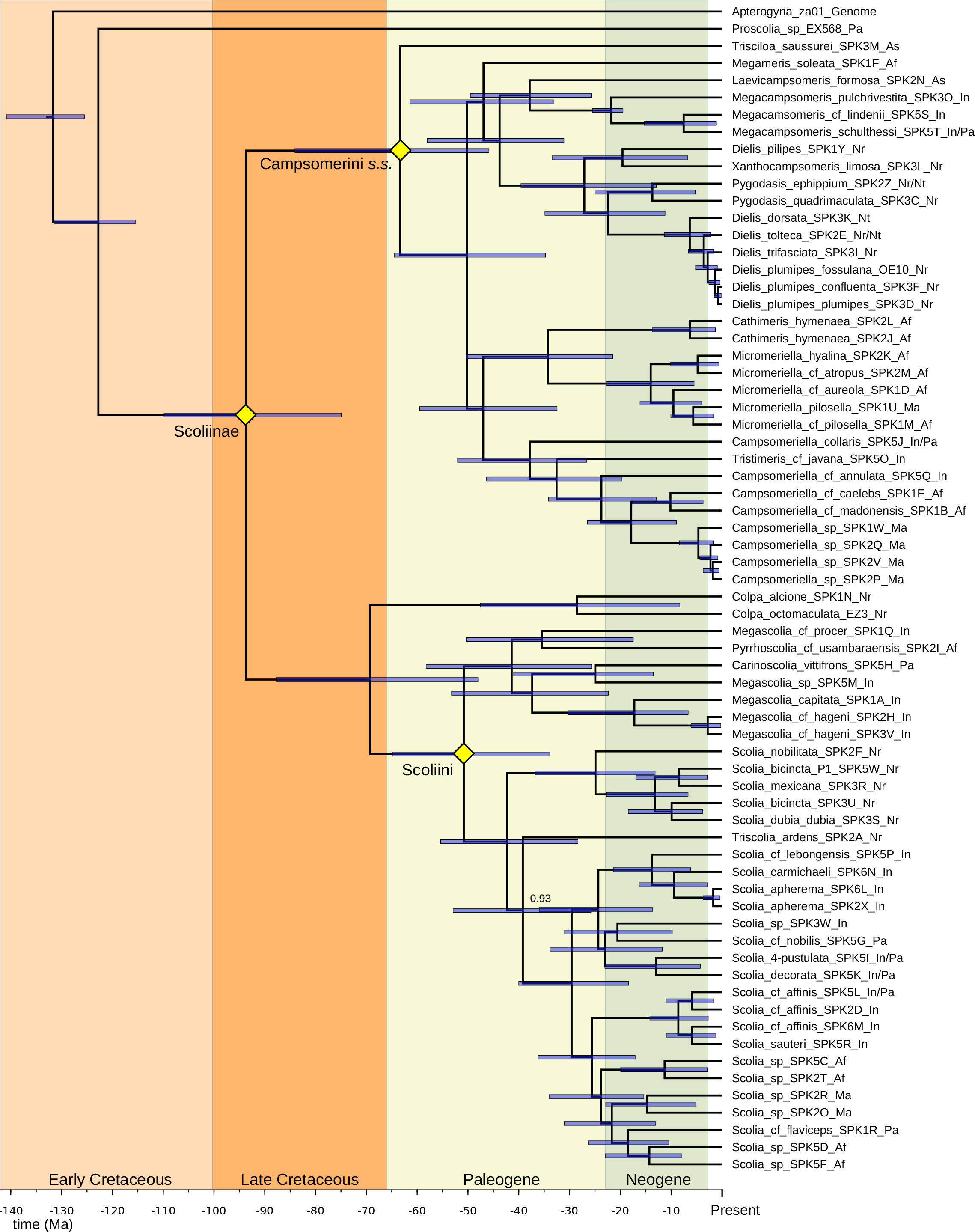
Bayesian MAP chronogram using additional calibrations on crown *Megacampsomeris* and crown Scoliidae (analysis 2b). All unlabeled internal nodes have posterior probability of 0.97 or greater. Taxonomic labels indicated with 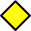.

There are some topological differences between the results of these analyses and the tree from analysis 2a, mostly in the relationships of Old World *Scolia* and the position of *Megameris soleata* as sister to (*Laevicampsomeris* + *Megacampsomeris*) + New World *Campsomerini sensu stricto*. However, both sets of analyses agree on the placement of *Colpa* as sister to the Scoliini and of *Triscolia ardens* as sister to the Old World *Scolia* clade.

#### Analysis 2c

The species or minimal clusters (SMC) tree inferred under the multispecies coalescent using STACEY (Fig. 11-12) recovered many of the same major clades as the other analyses. However, some relationships, particularly those that had conflicting resolutions among the previous analyses, were poorly resolved. Specifically, while *Colpa* is still sister to a monophyletic Scoliini and the *Megascolia* + *Pyrrhoscolia* + *Carinoscolia* clade is sister to all other Scoliini, the position of *Triscolia ardens* within the latter group is uncertain. *Trisciloa* is still sister to all other members of Campsomerini *sensu stricto*, the New World members of which form a monophyletic group. *Megameris soleata*, *Laevicampsomeris* + *Megacampsomeris*, and the New World Campsomerini *sensu stricto* form a clade, though the relationships among them is uncertain. Likewise, the relationships among this clade, the *Cathimeris* + *Micromeriella* clade, and the *Campsomeriella* + *Tristimeris* clade are not resolved.

**Figure 11.**
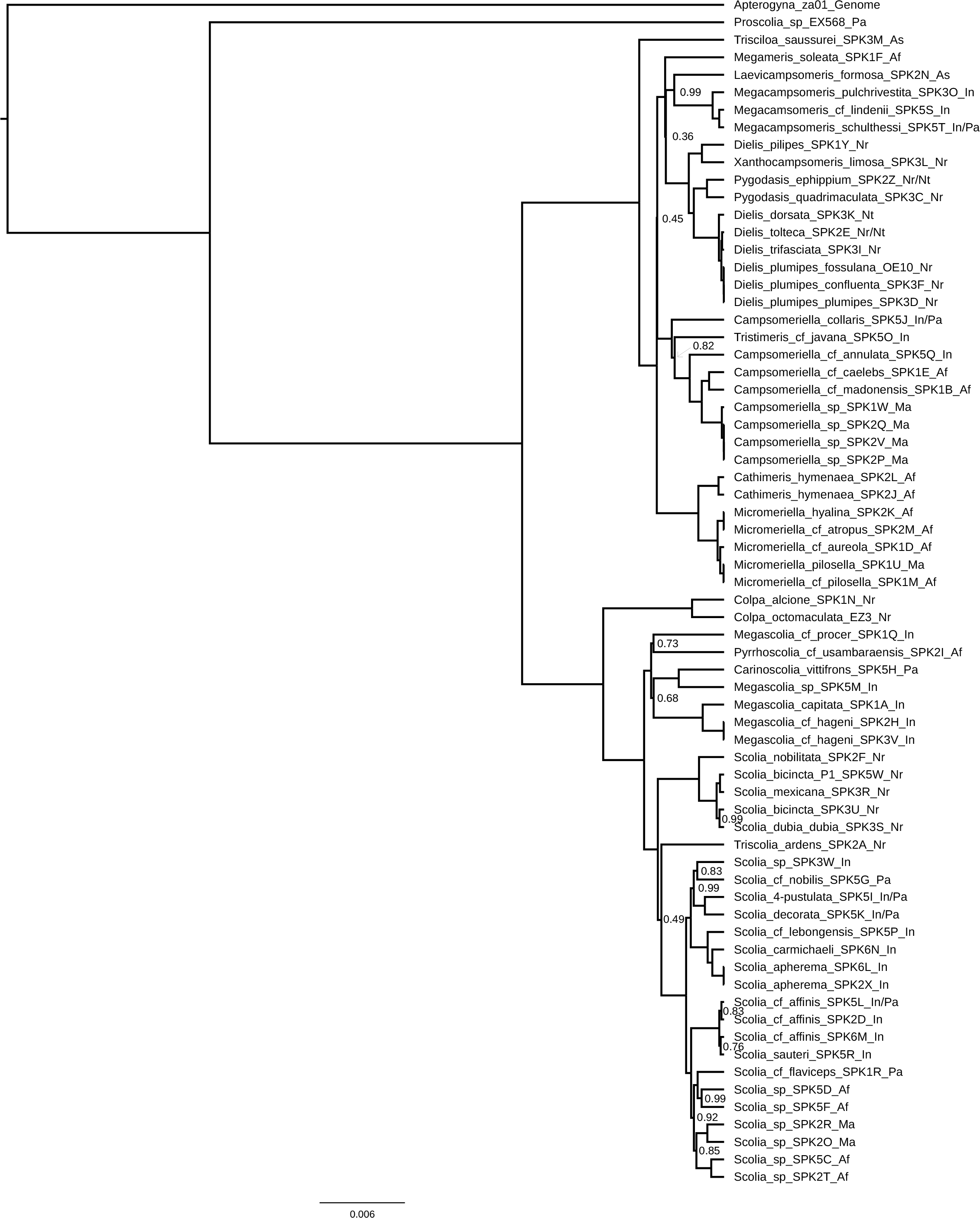
MCC species or minimal clusters tree based on 4 independent MCMC chains run using STACEY (analysis 2c). All unlabeled internal nodes have posterior probability of 1.0.

**Figure 12.**
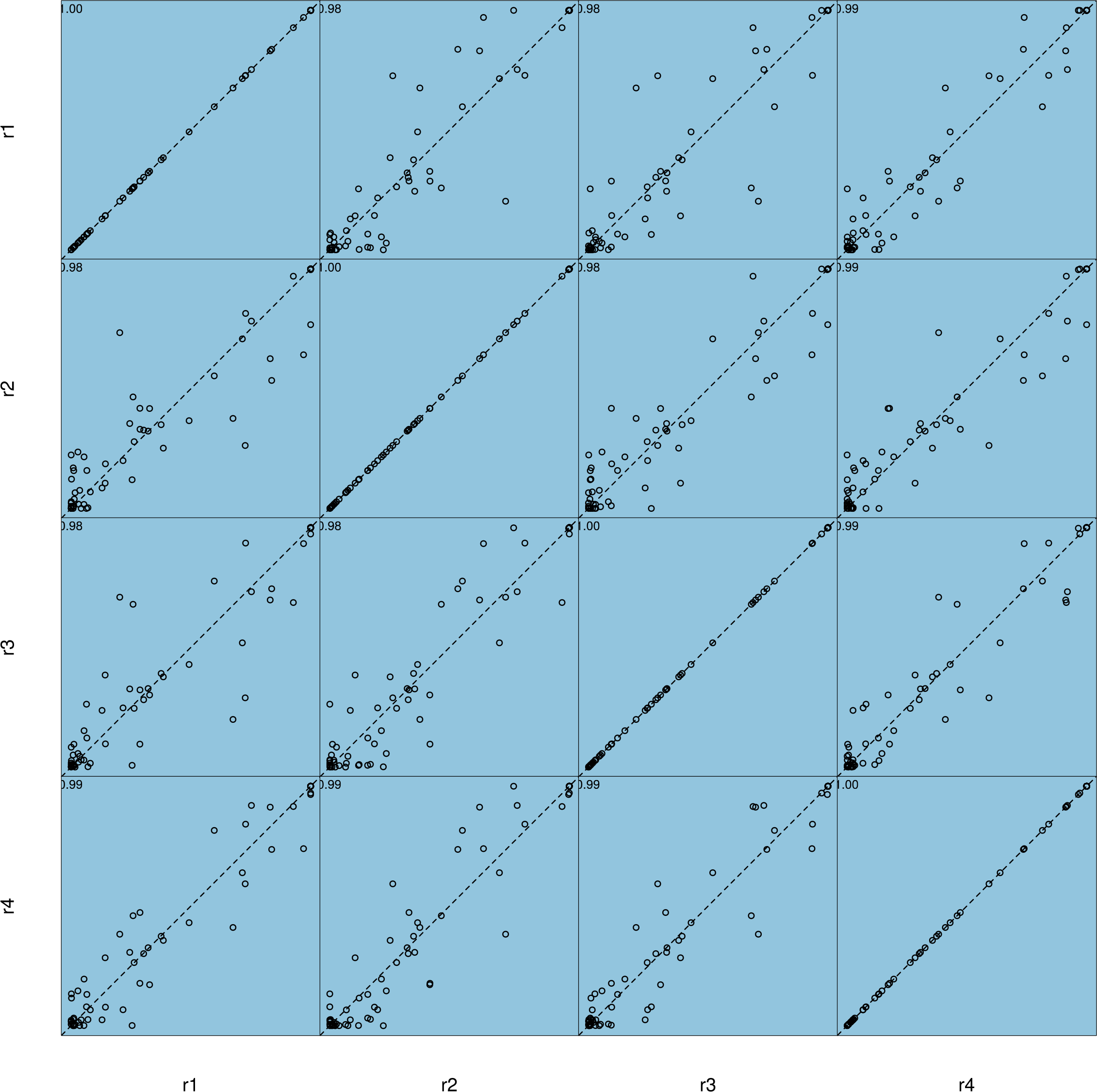
Comparison of split posterior probabilities between four independent MCMC runs using STACEY (analysis 2c). Numbers in the top left corners represent R^2^.

#### Analysis 2d

The “main ” ASTRAL topology, estimated using MCC trees only, was mostly congruent with the consensus topology, estimated using posterior samples and bootstrapping, in the case of the dataset with all loci (Fig. 13D) and of the dataset with loci having the highest-third combined posterior predictive effect sizes (Fig. 13C), with a few differences in the resolution of shallow nodes with low support. The dataset with loci having the lowest posterior predictive effect sizes showed somewhat bigger differences between the “main ” (Fig. 13A) and bootstrap consensus (Fig. 13B) topologies, the “main ” topology notably placing *Proscolia* as sister to the Campsomerini *sensus stricto*, albeit with low support.

**Figure 13.**
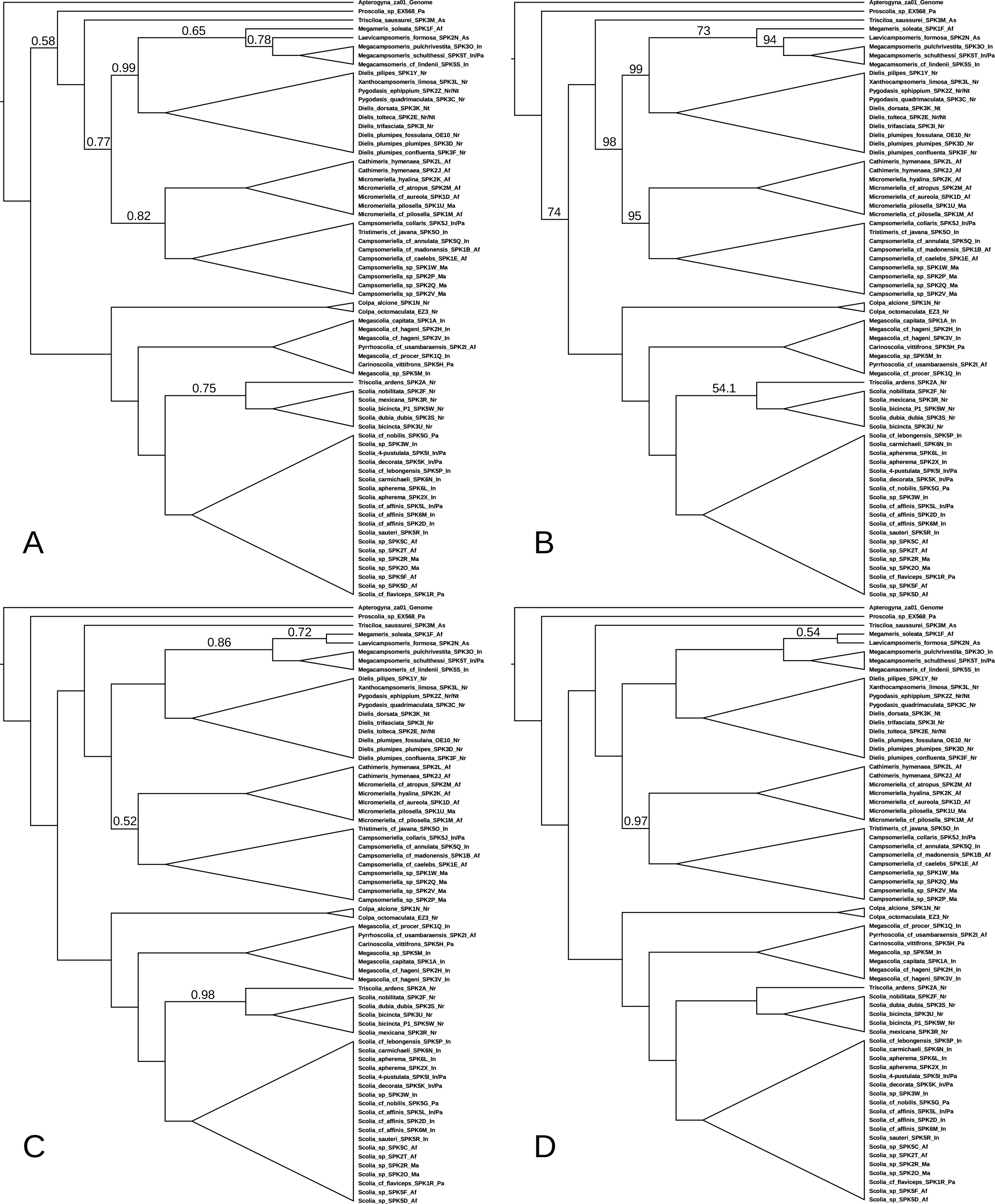
ASTRAL species trees (analysis 2d) based on MCC trees of (A) loci having the lowest (1/3) combined posterior predictive effect sizes, (C) loci having the highest (1/3) combined posterior predictive effect sizes, and (D) all loci. (B) is an ASTRAL bootstrap consensus tree using posterior samples of gene trees from loci having the lowest (1/3) combined posterior predictive effect sizes. Branch labels represent the local posterior probability or bootstrap support of/for the associated quadripartition.

The topology inferred using all loci mostly agrees with the results of analysis 2a, 2b, and 2c with the exception of *Megameris soleata* being inferred to be more closely related to *Laevicampsomeris* and *Megacampsomeris* than to the New World Campsomerini clade. Additionally, *Triscolia ardens* is placed as sister to the New World Scoliini, as opposed to being sister to the Old World *Scolia* clade as in analyses 2a and 2b and its position being unresolved as in analysis 2c. Analysis of the subset of loci with the highest combined posterior predictive effect sizes produced results almost identical to those based on all loci. Conversely, as reported above, using loci with the lowest posterior predictive effect sizes resulted in the unexpected placement of *Proscolia* as sister to Campsomerini *sensus stricto*. Relationships were otherwise similar to those inferred using other locus sets, but with lower local posterior probabilities associated with many quadripartitions.

#### Analyses 3a and 3b

Analyses 3a and 3b are based on data from 115 and 159 loci respectively. The results (Fig. 14-15) agree with each other and mostly agree with those from analysis 1b. Differences include *Triscolia ardens* being sister to the Old World scoliine clade and *Megameris soleata* being sister to *Laevicampsomeris formosa*.

**Figure 14.**
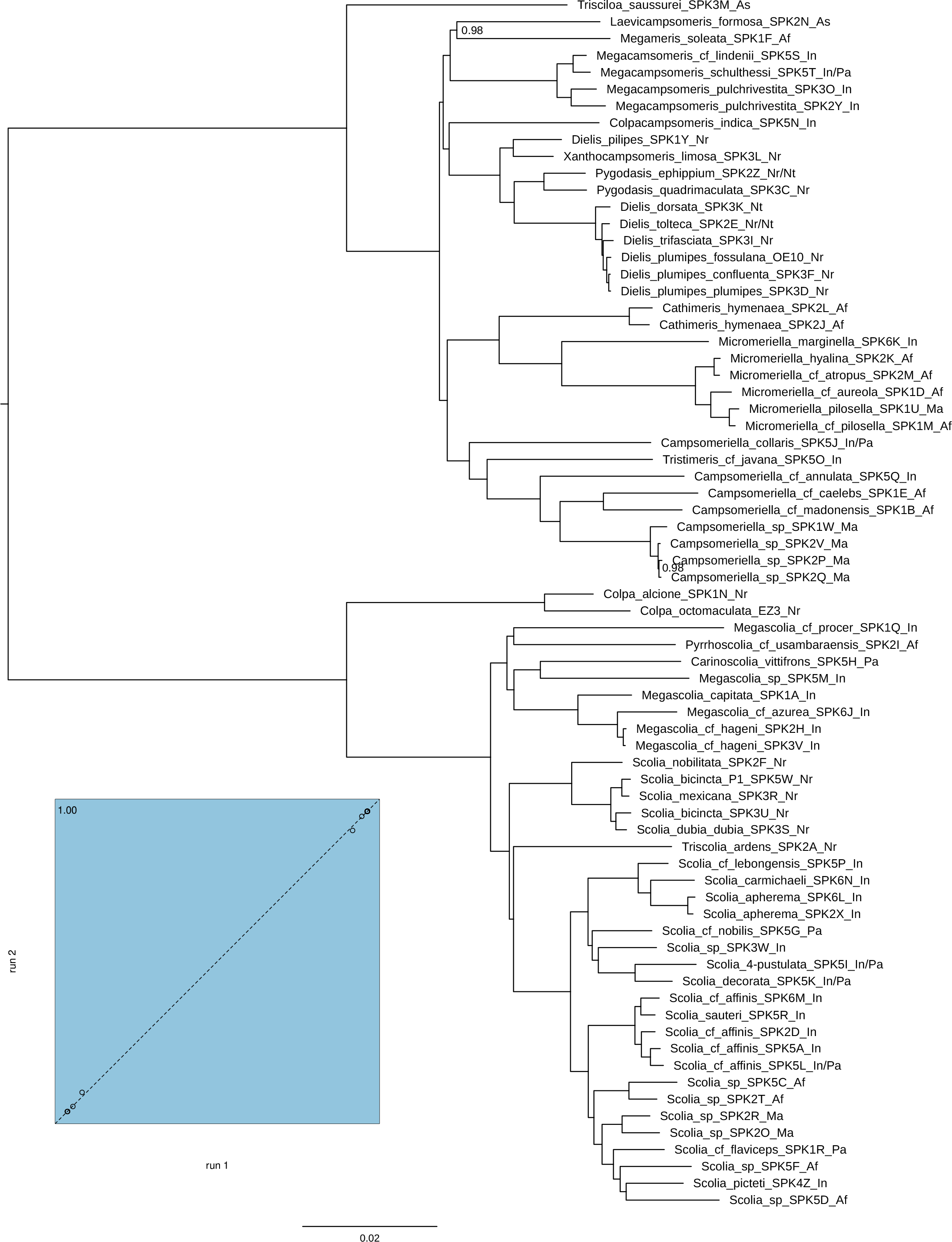
Bayesian MAP tree based on 115 UCE loci after data filtering using posterior predictive checks (analysis 3a). All unlabeled internal nodes have posterior probabilities of 1.0. Comparison of split posterior probabilities between two independent MCMC runs on lower left.

**Figure 15.**
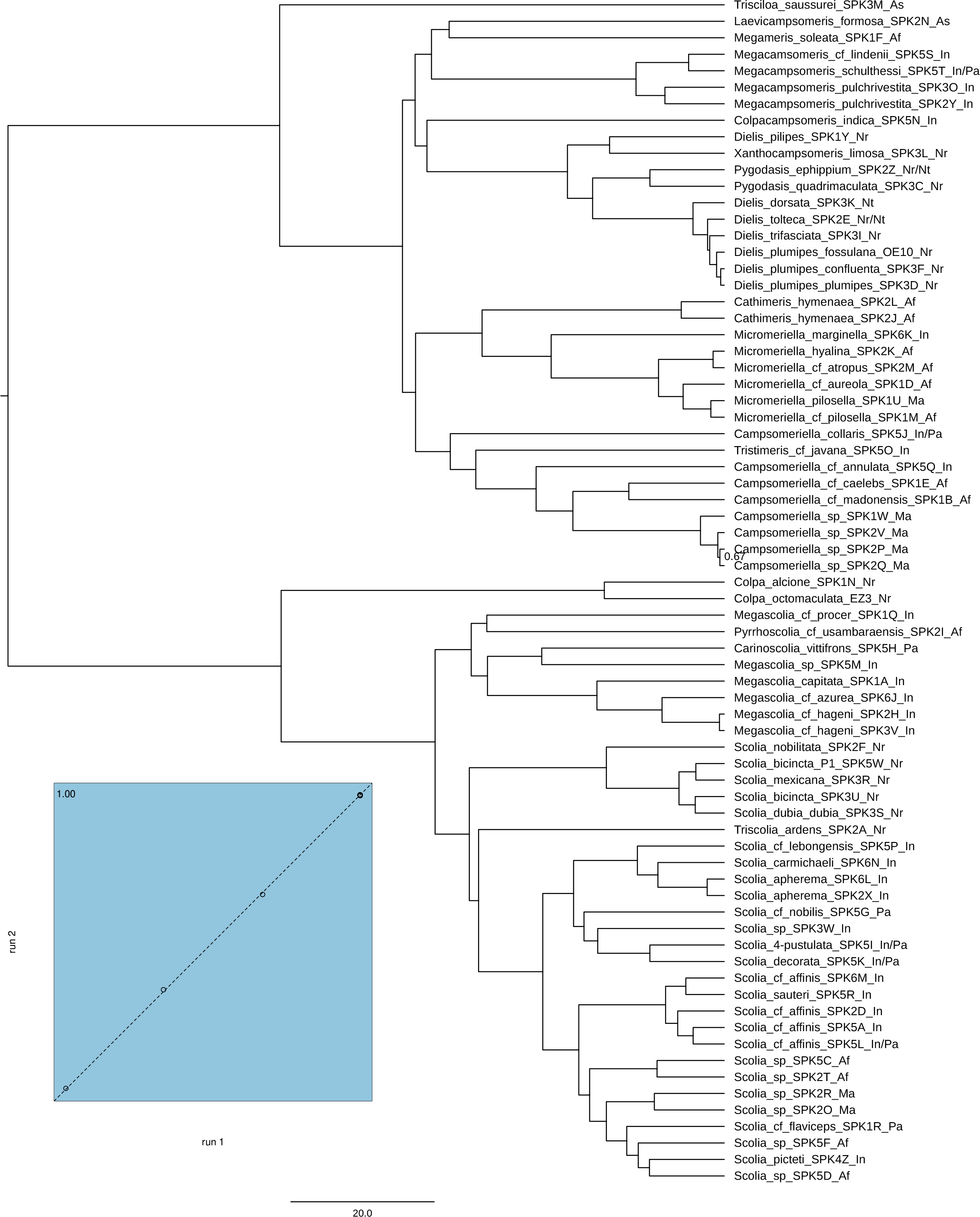
Bayesian MAP relative-time chronogram based on 159 UCE loci after data filtering using posterior predictive checks (analysis 3b). All unlabeled internal nodes have posterior probabilities of 1.0. Comparison of split posterior probabilities between two independent MCMC runs on lower left.

## Discussion

### Phylogenetic results and taxonomic implications

This is the first study to use molecular data to reconstruct the mammoth wasp phylogeny. Our results corroborate some long-standing phylogenetic hypotheses originally based on morphological data while contradicting others. Scoliid taxonomy has historically been unstable and confusing (see Elliott (2011) and Kimsey & Brothers (2016) for commentary). In the following discussion, we use Osten (2005) as the reference for the current status of taxon names unless otherwise specified. We use Campsomerini *sensu stricto* to refer to Campsomerini excluding *Colpa* and taxa more closely related to *Colpa* than to the Scoliini.

The genus *Proscolia* was originally described by Rasnitsyn (1977), hypothesized to be sister to the remaining extant Scoliidae, and placed in a new subfamily Proscoliinae, with the other extant Scoliidae relegated to the Scoliinae. Day *et al*. (1981) and Osten (2005) maintained this arrangement and treated the former subfamilies Scoliinae and Campsomerinae as the scoliine tribes Scoliini and Campsomerini respectively (Fig. 16C). Notable exceptions to this approach include earlier works by Osten (1988, 1993), where he argued against the inclusion of *Proscolia* in the Scoliidae, and Argaman (1996), who radically revised the higher-level scoliid taxonomy without conducting an explicit phylogenetic analysis. Argaman elevated the Campsomerini (minus *Colpa* and its presumed close relatives) back to subfamily rank (Fig. 16D) and placed it as sister to the remaining extant Scoliidae (including the Proscoliinae). Pilgrim *et al*. (2008) included three scoliids in their study and placed *Proscolia* as either sister to the other two scoliids or as sister to Bradynobaenidae + other Scoliidae. Two more recent molecular phylogenetic studies of aculeates that included five and three scoliid species respectively (Debevec *et al.,* 2012; Branstetter *et al.,* 2017a) placed *Proscolia* as sister to all other scoliids. All analyses in the present study (Fig. 16E) strongly support this placement.

**Figure 16.**
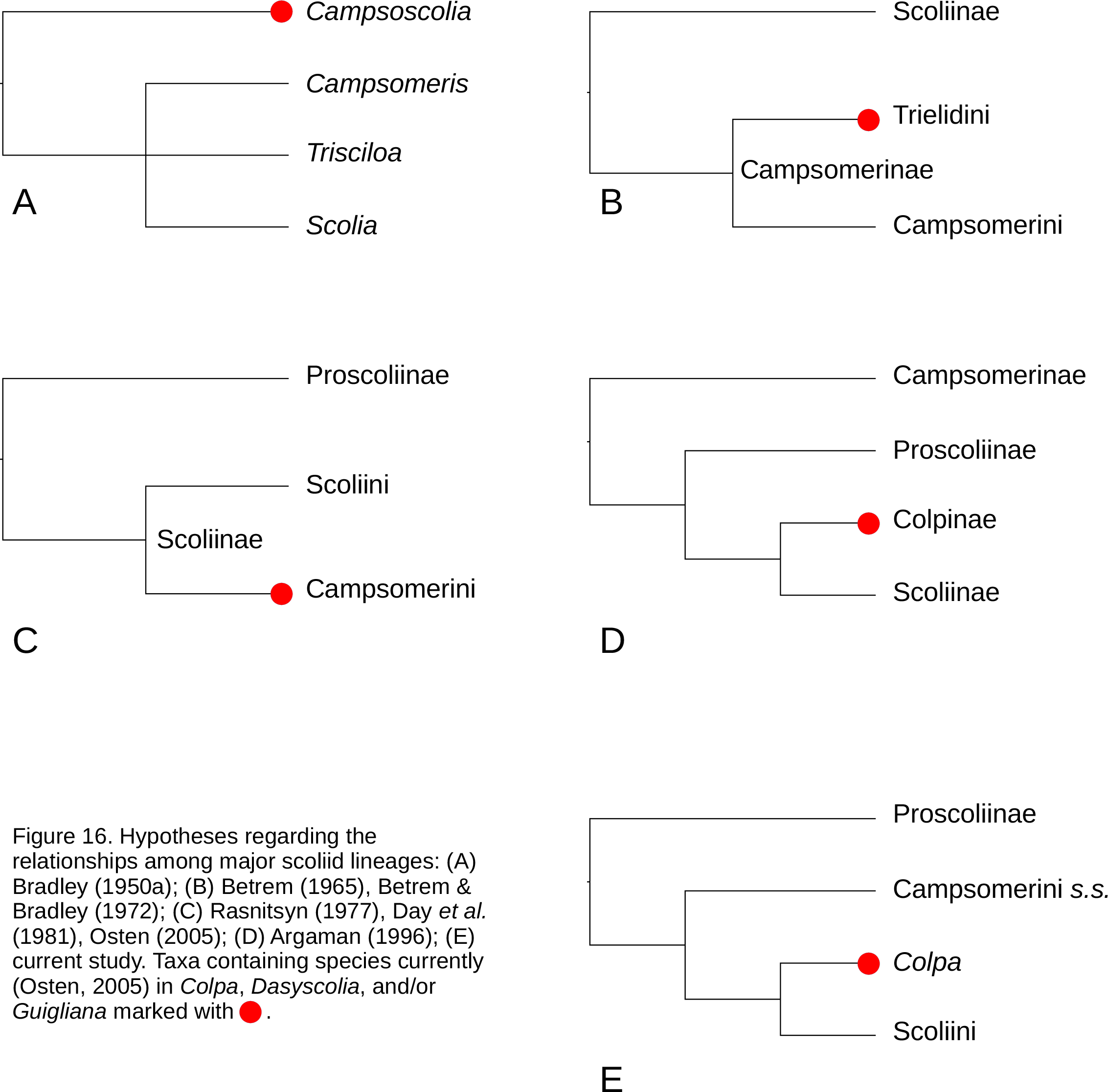
Hypotheses regarding the relationships among major scoliid lineages: (A) Bradley (1950a); (B) Betrem (1965), Betrem & Bradley (1972); (C) Rasnitsyn (1977), Day *et al*. (1981), Osten (2005); (D) Argaman (1996); (E) current study. Taxa containing species currently (Osten, 2005) in *Colpa*, *Dasyscolia*, and/or *Guigliana* marked with 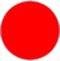.

The taxonomic treatment of the species currently comprising the genus *Colpa* has historically varied significantly. To date, none of the taxonomic changes have been supported by phylogenetic analyses. However, the following authors generally presented informal phylogenetic arguments when making taxonomic decisions. Bradley (1950a), using the name *Campsoscolia* for the genus including what is now *Colpa* and *Dasyscolia*, argued for a “basal ” placement of these taxa, presumably meaning they fall outside the clade formed by the remaining Scoliidae (Fig. 16A). Betrem (1965) erected the tribe Trielini (emended by Betrem & Bradley (1972) to Trielidini) within the Campsomerinae (Fig. 16B) to contain the genera *Trielis* (corresponding to *Campsoscolia* as used by Bradley (1950a) and currently understood (Day *et al*., 1981) to be a junior synonym of *Colpa*), *Crioscolia* (currently treated as a subgenus of *Colpa*), and *Guigliana*, which was formally described later by Bradley & Betrem (1967). Following the demotion of Campsomerinae to tribe rank by Day *et al*. (1981), *Colpa* and its allies were kept within the Campsomerini (Fig. 16C), with the implied relationships being Proscoliinae + (Campsomerini + Scoliini). Argaman (1996) on the other hand, created a new subfamily Colpinae (corresponding to the Trielidini of Betrem and Bradley (1972)) and placed it as sister to the Scoliini (which he elevated to subfamily rank), concluding that the Campsomerini *sensu stricto* (also elevated to subfamily rank) is sister to Proscoliinae + (Colpinae + Scoliinae) (Fig. 16D).

Debevec *et al*. (2012) included five scoliid species in their analyses, one of them being *Colpa sexmaculata,* but the main text contains no discussion of *Colpa* and the relationships within the Scoliidae. If we assume the monophyly of Campsomerini *sensu stricto* and of *Colpa* (each only represented by one species), the phylogenies included with the supporting information place Proscoliinae as sister to Campsomerini *sensu stricto* + (*Colpa* + Scoliini). All analyses in the current study agree with the latter hypothesis (Fig. 16E) while using a significantly larger dataset and attempting to mitigate the effects of non-randomly-distributed missing data and phylogenetic model violation.

In light of these results, morphological similarities between *Colpa* and the Campsomerini, such as the presence of an articulation between the basal and apical parts of the volsella and the presence of the second recurrent vein, are likely plesiomorphies. We recommend the exclusion of *Colpa* from Campsomerini when a formal taxonomic revision of Scoliidae is undertaken. However, a phylogenetic analysis establishing the positions of *Guigliana* and *Dasyscolia* (not represented in this study) should be considered a prerequisite of such a revision. Both genera lack the transverse impressed impunctate band on the frons, which serves as the defining feature of *Colpa*, but share with *Colpa* and the Scoliini some mesothoracic characters (Bradley, 1950a; Betrem & Bradley, 1972). If *Guigliana* and *Dasyscolia* form a monophyletic group with *Colpa*, the establishment of a tribe Colpini may be justified. Otherwise, if they are more closely related to or nested within the Scoliini, it may be reasonable to transfer *Colpa*, *Guigliana*, and *Dasyscolia* to that tribe. More complete sampling of this group would also allow the evaluation of its subgeneric classification. The subgenus *Colpa* (*Crioscolia*) has a strongly disjunct distribution in both the New and Old World (Bradley, 1950a). Our results (Fig. 2) indicate the paraphyly of *Colpa* (*Colpa*): the Nearctic *Colpa* (*Colpa*) *octomaculata* is more closely related to the Nearctic *Colpa* (*Crioscolia*) *alcione* than it is to the Palearctic *Colpa* (*Colpa*) *sexmaculata*. In addition to allowing a critical evaluation of the phylogenetic validity of *Colpa* subgenera, a molecular phylogeny including more *Colpa* species would contribute significant biogeographic information, as this group appears to have undergone dispersal and/or vicariance events between the Old World and the Americas independently of the Scoliini and the Campsomerini *sensu stricto*.

Campsomerini *sans Colpa* is inferred to be monophyletic in all our analyses, with *Trisciloa* always sister to the remaining members of the group. Likewise, all sampled New World Campsomerini *sensu stricto* form a clade with high support in all analyses. *Colpacampsomeris indica* is consistently inferred to be the closest relative of this New World clade in all analyses in which the former was included. However, we have not sampled any species from South America, so it remains unknown whether those share a closer relationship with the New World taxa sampled here or with Old World scoliids. *Dielis pilipes* groups with *Xanthocampsomeris* as opposed to with other *Dielis*. This is consistent with *D. pilipes* lacking some prominent morphological characteristics shared by other *Dielis*, such as a medial longitudinal furrow on the clypeus and a deep transverse furrow on the anterior of abdominal sternum II. Bradley (1964) states the opinion that *D. pilipes* should be excluded from *Dielis*, but this change was never formalized.

*Megacampsomeris* is always monophyletic in these analyses. Other consistently monophyletic groups include (1) *Micromeriella* with *Cathimeris* as the sister taxon and (2) the group consisting of *Campsomeriella*, *Tristimeris*, and some Malagasy species (undescribed or of uncertain taxonomic placement, provisionally labeled *Campsomeriella* sp. in the figures). Both *Tristimeris* and the Malagasy specimens are nested within *Campsomeriella*.

The positions of *Megameris* and *Laevicampsomeris* are uncertain, though they are likely more closely related to the New World Campsomerini, *Megacampsomeris*, and *Colpacampsomeris* than to other Campsomerini. In the current study, they are each represented by only one species. More thorough taxon sampling within these two genera will likely result in less uncertainty regarding their placement.

All analyses conducted here strongly support the monophyly of Scoliini. The first split within the Scoliini gives rise to two clades: one consisting of *Megascolia*, *Pyrrhoscolia*, and *Carinoscolia* and the other consisting of *Scolia* and *Triscolia*. *Megascolia* is consistently non-monophyletic in our analyses. The situation warrants a taxonomic revision, though it should ideally be informed by future phylogenetic studies that are able to sample *Megascolia*, *Pyrrhoscolia*, and *Carinoscolia* more completely. Sequencing of multiple *Carinoscolia* species is especially important, given that the genus is suggested to be polyphyletic by Golfetti (2019).

Our sampling of New World species was restricted to the Nearctic, and the affinities of Neotropical scoliines thus remain uncertain. However, all sampled Nearctic *Scolia* form a single clade. The phylogenetic position of *Triscolia ardens* was inconsistent across our analyses. The genus *Triscolia* has a complicated taxonomic history (see Betrem & Bradley, 1964) and currently includes only two Nearctic species, *T. badia* and *T. ardens*. In all phylogenies where *Scolia verticalis* is included, *T. ardens* and *S. verticalis* are sisters. This is somewhat surprising given that *S. verticalis* is an Australasian species. We have mostly ruled out contamination and misidentification (see results section above) as potential explanations. More thorough sampling of scoliines from Australasia, Southeast Asia, and the eastern Palearctic might reveal species related to *S. verticalis* and fill in the gap in distributions, making a relationship with the Nearctic fauna more plausible. It is also possible that the two species of *Triscolia* are the only extant representatives of a previously more widespread lineage. The lack of close relatives of either species in the present study also means they are both subtended by long branches. A combination of the potential for long branch attraction and the disproportionately high fraction of missing data from *S. verticalis* raises the suspicion that the pairing might be artefactual. Regardless of its relationship to *S. verticalis*, *T. ardens* is recovered in our analyses either as closely related to the Nearctic *Scolia* clade or to the Old World *Scolia* clade, making it likely that the genus *Scolia* is paraphyletic irrespective of which placement of *T. ardens* is correct. One potential course of action is to synonymize *Triscolia* with *Scolia*. However, any taxonomic decisions involving *Scolia* should take into account the phylogenetic positions of two other large Scoliine genera, *Liacos* and *Austroscolia*, both of which are not represented in the current study.

Given the proliferation of scoliid generic names attached to groups defined mainly by superficial characters such as color and punctation, it seems likely that there are many examples of distinctive groups within larger genera being given their own generic names, thus rendering the larger genera paraphyletic. Further phylogenetic studies with more complete taxon sampling are needed before a taxonomic revision of scoliid genera is attempted. In the absence of such studies, we recommend proceeding cautiously when describing new species (such as those belonging to the Malagasy scoliid fauna) and avoiding the establishment of new genera or groups of higher rank without first conducting thorough phylogenetic analyses.

### Divergence times and biogeography

The precision of node age estimates in the current study is limited by the small number of fossils that can be reliably attributed to the scoliid crown. It might be possible to slightly increase precision by conducting analyses with a broader phylogenetic scope. Including taxa from the Apoidea and Formicoidea could allow fossil data from those clades to inform overall rates of molecular evolution. However, apoids and formicoids being much more diverse than scoliids makes it difficult to sample species evenly across clades, and care must be taken to accommodate for this in any attempted analyses. The increased likelihood of heterogeneity in the evolutionary process becoming problematic as one expands the scope of the analysis should also be considered and addressed. Ultimately, the discovery and description of well-preserved crown fossils is likely to be a necessary prerequisite to achieving scoliid divergence time estimates with better precision and accuracy.

Due to weak sampling from some biogeographic regions, particularly Australasia and the Neotropics, we did not conduct a formal phylogeographic analysis. However, our phylogenetic results do indicate some biogeographic patterns that could be further investigated in future studies.

We estimated the stem age of the Nearctic Campsomerini *sensu stricto* clade to be between 19 Ma and 46 Ma (95% HPD interval) when calibrating the root age only (Fig. 8). Among taxa sampled in this study, the closest relatives of this clade are taxa from Indomalaya, Australasia, and the eastern Palearctic. This suggests a possible exchange of fauna across Beringia during the Oligocene or later Eocene, which is broadly consistent with patterns observed in other animal groups (Jiang *et al*., 2019). The Nearctic *Scolia* clade has a very similar estimated stem age (19-50 Ma). Analyses using additional (but less reliable, in terms of the phylogenetic placement of the associated fossil) calibrations extend the age 95% credible intervals into the early Eocene. Further refinement of node age estimates, in conjunction with more complete geographic sampling, is needed to evaluate the possibility of late (*c*. 65 Ma) exposures of the Thulean Route (Brikiatis, 2014) contributing to scoliid dispersal.

The phylogenetic position of *Triscolia* is uncertain, and has implications for the number and timing of biotic interchanges between North America and other regions. In addition to resolving the position of *Triscolia*, future phylogenetic studies need to prioritize sampling of the South American and Australasian scoliids. It is currently unclear whether South American Scoliini and Campsomerini *sensu stricto* each represent single lineages or multiple lineages with different biogeographic origins. It is possible that South America harbors relatively young lineages originating from Africa or the Nearctic and dispersing into South America during the Late-Early Eocene or later (Hoffmeister, 2020) and/or more ancient lineages with possible relationships to the Australian and African fauna. Understanding the phylogenetic and biogeographic affinities of South American scoliids, while interesting in itself, is also essential to understanding patterns of scoliid diversification and answering questions such as why the Campsomerini *sensu stricto* are significantly more diverse than the Scoliini in the New World tropics while the opposite pattern holds in the Afrotropic and Indomalaya (Bradley, 1950b; 1959).

Madagascar is home to members of at least two campsomerine lineages, represented in the current study by one species of *Micromeriella* and several samples (probably from currently undescribed species) falling within the *Campsomeriella* clade. The presence of *M. pilosella* is probably due to a very recent dispersal from mainland Africa, while the Malagasy *Campsomeriella* lineage is older but also most closely related to African species. Given that the Malagasy scoliid fauna has received much less study than that of mainland Africa, it is certainly possible that among the species not sampled in this study there exist representatives of older endemic lineages that are not closely related to either *Micromeriella* or *Campsomeriella*. Our study additionally included two (probably undescribed) species belonging to *Scolia*. The scoliine genera *Liacos* and *Autroscolia* both have representatives on the African mainland, Madagascar, Asia, and Australia (Bradley, 1950b; Osten, 2005; Elliott 2011), while the morphologically distinctive *Mutilloscolia* is confined to Madagascar (Bradley, 1959). None are included in this study and their phylogenetic relationships to other scoliines remain poorly understood. Although it is possible that these genera could be nested within *Scolia* (which is mostly identified by lacking the defining characters of other genera), the apparent lack of morphological characters uniting them specifically with the *Scolia* species sampled here suggests that the current Malagasy scoliine diversity is likely a result of multiple dispersal (and possibly vicariance) events. This is tentatively supported by the morphology-based phylogenies of Golfetti (2019), which place *Austroscolia* and *Liacos* outside the clade formed by all scoliini sampled in this study.

### Methodological considerations

Doyle *et al*. (2015) demonstrated the potential utility of filtering data using posterior predictive methods. We made use of a similar approach, albeit limiting it to data-based (Huelsenbeck *et al*., 2001) as opposed to inference-based (Brown, 2014a) tests. Molloy & Warnow (2018) used a simulation-based approach to explore the effect of data filtering using various criteria on species tree inference using ASTRAL (among other methods). They found that excluding loci with high gene tree estimation error can improve the accuracy of species tree inference when levels of incomplete lineage sorting (ILS) were moderate to low. The dependence on ILS levels was explained in terms of the number of gene trees required to accurately reconstruct the species tree increasing with higher levels of ILS. Thus, the negative effect of using fewer genes sometimes outweighed the positive effect of more accurate gene trees (Molloy & Warnow, 2018). In this context, we make the following observations based on our empirical analyses:

Using posterior predictive p-values with “conventional ” cutoffs (e.g. 0.05) resulted in the exclusion of the majority of available loci. In some cases (e.g. Fig. 13A), this led to an unexpected and implausible species tree topology resulting from ASTRAL analyses (i.e. placement of *Proscolia* as sister to the Campsomerini). This could be a result of too few loci being used. Additionally, one would expect a correlation between the amount of data and the ability to detect model inadequacy, which might lead to the retention of less “informative ” loci. This appears to be borne out in analysis 2d, where mean pairwise Robinson-Foulds distances among posterior topology samples were on average higher (73.3 versus 53.8) for the third of loci having the lowest posterior predictive effect sizes compared to the third having the highest. Under these circumstances, a fully-resolved point estimate of the topology might be a worse representation of the gene tree posterior distribution, and variance among gene tree point estimates might be higher, even if there is no ILS and the underlying posterior distributions are unbiased. This could explain why we observed generally lower quadripartition support values resulting from analyses of loci with lower posterior predictive effect sizes even when the number of loci per analysis was kept constant (Fig. 5A, B; Fig. 13A, C). In contrast to Fig. 13A, a bootstrap-based ASTRAL species tree (Fig. 13B) that used samples from the posterior distributions of gene trees (as opposed to point estimates) recovered *Proscolia* in a more plausible position that is also corroborated by our STACEY analysis. Mirarab (2019) observed that using samples from gene tree posteriors does not have the same negative effect on species tree accuracy as does using gene tree bootstrap replicates in a ML framework. We concur with Mirarab (2019) that further investigation is warranted. Potential use cases for this hybrid approach could be datasets with both (1) a limited number of genes available (e.g. from Sanger data) where the accuracy of estimates using ASTRAL with gene-tree point estimates may be lower and (2) with a very large number of terminal taxa where a fully Bayesian approach (model-based coestimation of gene trees and species tree) may be more computationally challenging.

## Data availability

Raw sequences are available at: TBD

Other data (assemblies, multiple sequence alignments, etc.) and raw output files that support the findings of this study are available at: TBD

Scripts used for analyses are available at: TBD

## Acknowledgments

We would like to thank M. Hauser (California Department of Food and Agriculture), S.L. Heydon (Bohart Museum of Entomology, University of California, Davis), and K. Williams (California Department of Food and Agriculture) for providing access to scoliid specimens at their respective institutions; B.E. Boudinot, M. Hauser, C. Parker, and T. Zavortink for giving access to specimens in their personal collections; former members of the Ward Ant Lab, University of California, Davis (UC Davis) M. Borowiec, B.E. Boudinot, and M. Prebus and A. Abrieux from the Chiu Lab (UC Davis) for training the lead author in wet lab techniques; B. Moore (UC Davis) for mentoring the lead author in phylogenetic methods; G. Attardo, J. Bond, J. Chiu, B. Johnson, S. Nadler, and P.S. Ward for the use of their respective labs and equipment at UC Davis; L. Smith (Evolutionary Genetics Lab, Museum of Vertebrate Zoology, University of California, Berkeley) for providing access to a sonicator and training the lead author in its use; B. Boudinot and C. Pagan (Nadler Lab, UC Davis) for assistance with library preparation and enrichment; former FARM HPC cluster sysadmins B. Broadley and T. Thatcher for help with using the cluster; current and former members of the Ward Ant Lab M. Borowiec, B.E. Boudinot, Z. Griebenow, Z. Lieberman, J. Oberski, M. Prebus, and P.S. Ward and of the Moore Lab (UC Davis) E. Espejo, J. Gao, M.R. May, and B. Moore for helpful and insightful discussion; N. Tam for proofreading and providing comments on the manuscript.

