## Supplementary material for "The evolutionary history of mammoth wasps (Hymenoptera: Scoliidae)": Figure_S2_T.ardens_and_S.verticalis_validation.pdf (from another version of this preprint: version 1, 2022-01-27)

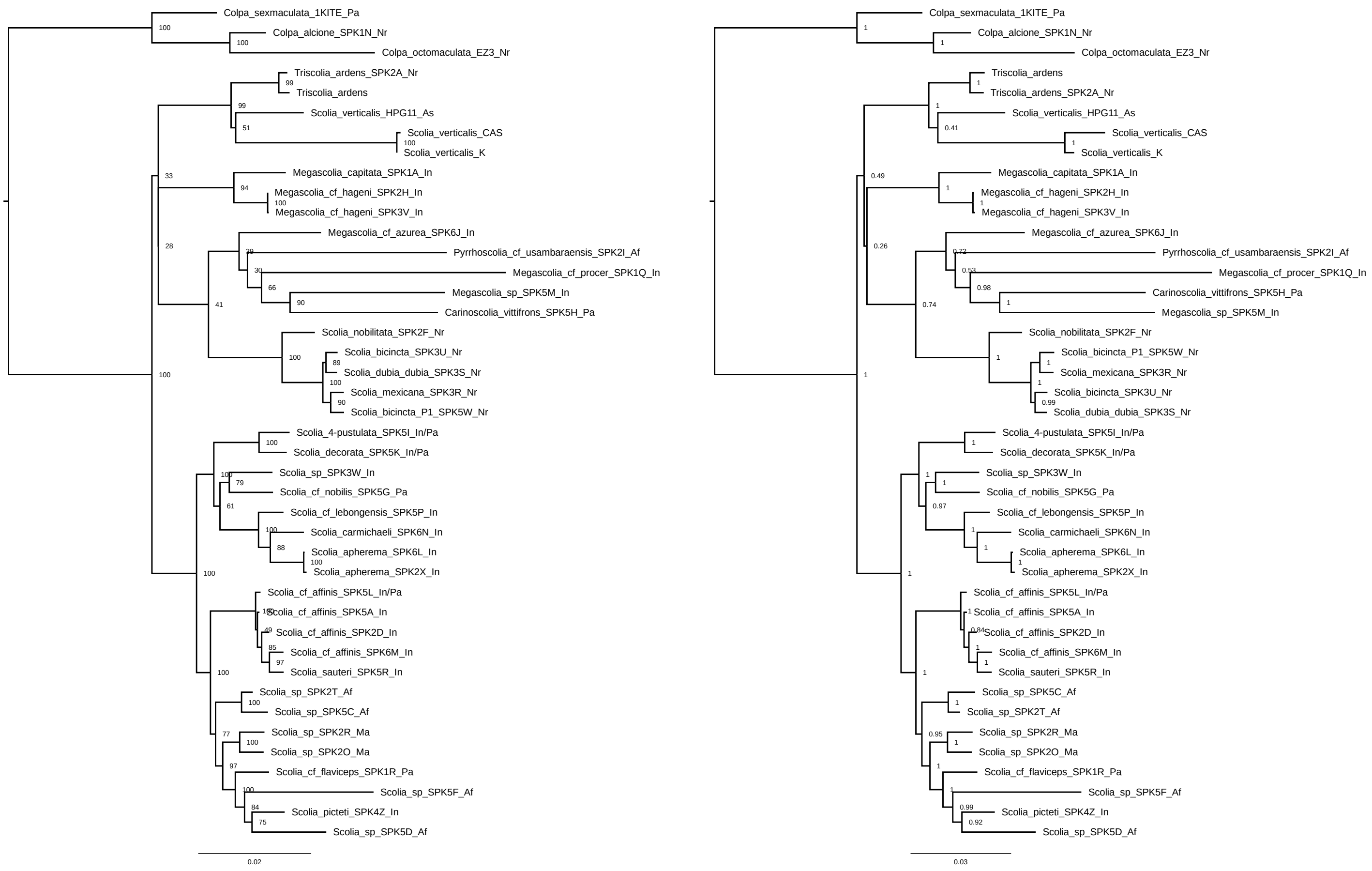


Figure S2. (A) Maximum likelihood tree; node labels are ultrafast bootstrap support. (B) Bayesian MAP tree; node labels are posterior probabilities.
